# Serum Amyloid A Proteins Induce Pathogenic T_H_17 Cells and Promote Inflammatory Disease

**DOI:** 10.1101/681346

**Authors:** June-Yong Lee, Jason A. Hall, Lina Kroehling, Lin Wu, Tariq Najar, Henry H. Nguyen, Woan-Yu Lin, Stephen T. Yeung, Hernandez Moura Silva, Dayi Li, Ashley Hine, P’ng Loke, David Hudesman, Jerome C. Martin, Ephraim Kenigsberg, Miriam Merad, Kamal M. Khanna, Dan R. Littman

**Author notes:** Contributed equally (June-Yong Lee, Jason A. Hall).

## Abstract

Lymphoid cells that produce IL-17 cytokines protect barrier tissues from pathogenic microbes, but are also prominent effectors of inflammation and autoimmune disease. T-helper (T_H_17) cells, defined by RORγt-dependent production of IL-17A and IL-17F, exert homeostatic functions in the gut upon microbiota-directed differentiation from naïve CD4^+^ T cells. In the non-pathogenic setting, their cytokine production is regulated by serum amyloid A proteins (SAA1 and SAA2) secreted by adjacent intestinal epithelial cells. However, T_H_17 cell behaviors vary markedly according to their environment. Here we show that SAAs additionally direct a pathogenic pro-inflammatory T_H_17 cell differentiation program, acting directly on T cells in collaboration with STAT3-activating cytokines. Using loss- and gain-of-function mouse models, we show that SAA1, SAA2, and SAA3 have distinct systemic and local functions in promoting T_H_17-mediated inflammatory diseases. These studies suggest that T cell signaling pathways modulated by the SAAs may be attractive targets for anti-inflammatory therapies.

## Introduction

T-helper 17 (T_H_17) cells and related IL-17-producing T cells perform critical roles at mucosal surfaces, mediating protection from pathogenic bacteria and fungi and contributing to regulation of the mutualistic organisms that comprise the microbiota (Honda and Littman, 2016). T_H_17 cells also drive the pathogenesis of multiple inflammatory diseases (McGeachy et al., 2019; Patel and Kuchroo, 2015; Stockinger and Omenetti, 2017). Yet, despite extensive studies conducted both *in vitro* and *in vivo*, the differentiation cues that distinguish T_H_17 cells that execute homeostatic functions, such as maintenance of barrier epithelial integrity, from those with potentially harmful inflammatory functions as observed in autoimmune diseases, remain an enigma. Differentiation of the latter, which have a distinct transcriptional program with features resembling those of T_H_1 cells, requires IL-23 and IL-1β (Chung et al., 2009; Hirota et al., 2011; Komuczki et al., 2019; McGeachy et al., 2009). Mice deficient for either of those cytokines retain the capacity to induce commensal microbe-specific T_H_17 cells (Ivanov et al., 2008), but are largely resistant to autoimmune disease in multiple animal models (Kullberg et al., 2006; Langrish et al., 2005). *In vitro* differentiation of the “homeostatic” T_H_17 cells can be achieved by stimulation of antigen-activated T cells with TGF-β and either IL-6 or IL-21 that induce STAT3 phosphorylation, which is prerequisite for expression of the T_H_17-defining transcription factor, RORγt (Ivanov et al., 2006; Zhou et al., 2007). When myelin-reactive T cells were generated *in vitro* with TGF-β and IL-6, they failed to provoke autoimmune disease following transfer into mice (Lee et al., 2012). However, additional *in vitro* exposure to IL-23 rendered these cells pathogenic, as did their differentiation in the absence of TGF-β, with a combination of IL-6, IL-1β, and IL-23 (Ghoreschi et al., 2010; Lee et al., 2012).

The SAAs are a family of acute phase response proteins that were recently associated with gut microbial ecology and inflammation and are encoded by at least four closely-linked genes that likely arose from gene duplication events (Lloyd-Price et al., 2019; Tang et al., 2017; Uhlar et al., 1994). SAA1,2, and 3 are elicited by inflammatory cues, while SAA4 is constitutively produced and regulated independently of the inflammatory state (De Buck et al., 2016; Yarur et al., 2017; Ye and Sun, 2015). Previously, we demonstrated that colonization of mice with an epithelial cell associated commensal microbe, segmented filamentous bacteria (SFB), triggered local secretion of SAA1 and SAA2 by the IEC. The SAAs directly potentiated local T_H_17 cell effector cell function (Ivanov et al., 2009; Sano et al., 2015). In this context, SAA1/2 remained confined to the ileum and did not enter into the circulation (Sano et al., 2015). However, SAA1 and SAA2 are most prominently induced upon liver injury and/or inflammation, with concentrations in serum increased by as much as a thousand-fold over basal levels (De Buck et al., 2016; Yarur et al., 2017; Ye and Sun, 2015). Serum concentrations of SAA1 and SAA2 are also consistently elevated in various T_H_17-mediated autoimmune diseases, including rheumatoid arthritis, Crohn’s disease (CD), ulcerative colitis (UC), and multiple sclerosis, in addition to numerous cancers (Ye and Sun, 2015). Systemic SAA levels have long served as biomarkers, but whether they contribute to these diseases is not known. The secreted proteins form hexamers that associate in serum with high density lipoprotein, and are thought to be involved in the maintenance of lipid homeostasis (Wang and Colon, 2004; Wang et al., 2002). There is also evidence that SAAs, particularly SAA3, may form tetramers and associate with derivatives of vitamin A (Derebe et al., 2014). The anatomical distribution and cellular targets of SAAs in chronic disease are largely undefined, but the SAAs were recently implicated in conditioning the liver microenvironment to provide a niche for tumor metastases (Lee et al., 2019).

Here, we investigate the role of SAAs in the differentiation and function of potentially pathogenic T_H_17 cells. We find that the SAAs can substitute for TGF-β in the induction of Th17 cells, but engage a distinct signaling pathway that results in a pro-inflammatory program of differentiation. As a consequence, SAAs contribute in vivo to T_H_17-mediated pathogenesis, revealed in inflammatory bowel disease and experimental autoimmune encephalomyelitis with both loss- and gain-of-function models. Collectively, our findings show that the SAAs contribute selectively to T_H_17 cell functions in vivo, and suggest strategies for therapeutic modulation in T_H_17-mediated inflammatory disease.

## Results

### The SAAs direct a distinct T_H_17 differentiation program independently of TGF-β

*In vitro* activation of naïve murine CD4^+^ T cells in the presence of IL-6 alone is insufficient to promote T_H_17 cell differentiation (Bettelli et al., 2006; Mangan et al., 2006; Veldhoen et al., 2006). Addition of either TGF-β or IL-1β plus IL-23 is required for the T cells to up-regulate RORγt and produce T_H_17 cytokines. Unexpectedly, we observed that a combination of SAA1 and IL-6 (referred to T_H_17-SAA1 condition), in the presence of TGF-β neutralizing antibody, elicited potent T_H_17 cell differentiation in a dose-dependent manner, with expression of the signature cytokines, IL-17A and IL-17F (Figures 1A-C and Figure S1A). Similar results were obtained using T cells expressing the dominant negative form of TGFβRII (TGFβRII DNtg), while addition of SAA1 neutralizing antibodies prevented T_H_17 cell induction under T_H_17-SAA1 conditions (Figures S1B and S1C). Expression of RORγt was similar between T_H_17-SAA1 cells and T_H_17 cells induced by the standard polarization cocktail consisting of IL-6 + TGF-β (T_H_17-TGFβ) (Figure 1D). Notably, naïve murine CD8^+^ T cells responded similarly to CD4^+^ cells under the different polarization conditions (Figures S1D and S1E). To determine whether SAA-directed signaling converged with the TGF-β signaling pathway, we assessed levels of phosphorylated SMAD2/3 (pSMAD2/3). As expected, TGF-β rapidly induced pSMAD2/3. However, this was not observed with SAA1, which instead was found to rapidly engage the MAPK pathway, inducing p38 phosphorylation (Figure 1E and S1F). These findings demonstrate that SAA1 promotes Th17 cell differentiation through a TGF-β−independent mechanism.

**Figure 1.**
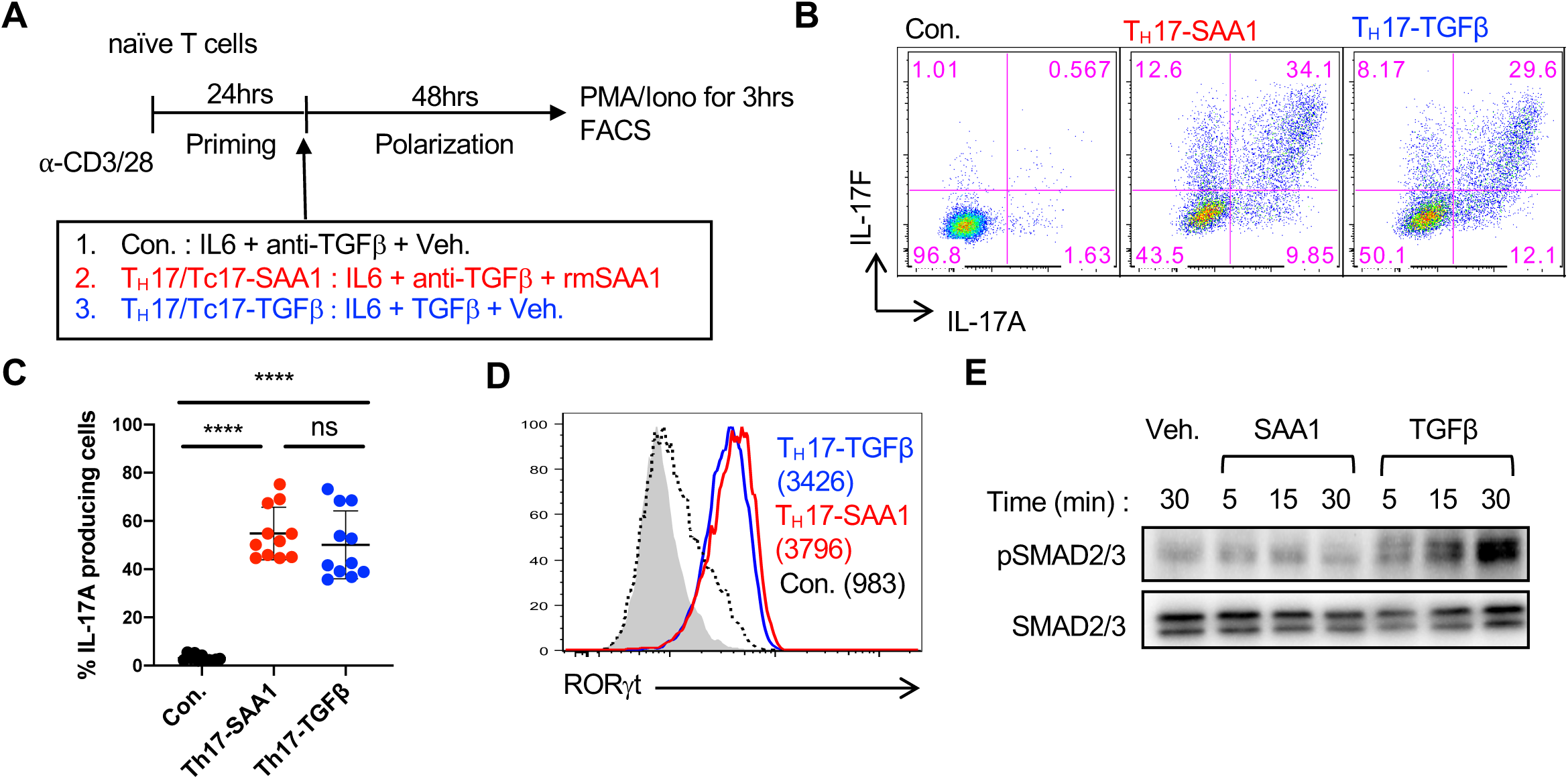
SAAs act directly on mouse T cells to induce T_H_17 cell differentiation *in vitro* in absence of TGF-β. **(A)** Experimental scheme for *in vitro* differentiation of naïve CD4^+^ or CD8^+^ T cells. **(B and C)** Flow cytometric analysis of IL-17A and IL-17F expression (**B**) and summary of IL-17A frequency (**C**) among re-stimulated cells. Summary of 3 experiments, with n = 11. Statistics were calculated using the unpaired two-sided Welch’s t-test. Error bars denote the mean ± s.d. ns = not significant, ****p < 0.0001. **(D)** RORγt expression in T_H_17 cells. Geometric mean fluorescence intensities (gMFI) are included in parentheses. Representative data of n > 10 experiments. **(E)** Immunoblotting for SMAD2/3 phosphorylation (pSMAD2/3) of primed CD4^+^ T cells upon rmSAA1 (10μg/ml) or TGF-β (1ng/ml) treatment for indicated times. Total SMAD2/3 is shown as a loading control. See also Figure **S1**.

To compare *in vitro* T_H_17 cell programming by SAA1 and TGF-β, we performed RNA sequencing (RNAseq) of cells differentiated under T_H_17-SAA1 or T_H_17-TGFβ conditions for 3h, 12h, and 48h (Figure S2A). After 48h, there were 3537 differentially expressed (DE) genes between T_H_17-SAA1 and T_H_17-TGFβ cells (Figure 2A). Compared to T_H_ cells cultured in IL-6 alone or the T_H_17-TGFβ condition, T_H_17-SAA1 cells exhibited a profound induction of hallmark chronic inflammatory disease-associated genes, such as *Il23r* (Abdollahi et al., 2016; Duerr et al., 2006; Gaffen et al., 2014; Hue et al., 2006), *Il1r1* (Shouval et al., 2016), and S*100a4* (Oslejskova et al., 2009) (Figure S2B and S2C). Gene set enrichment analysis using previously defined T_H_17 datasets revealed a striking correlation with the “pathogenic” T_H_17 signature, but an anti-correlation with the “non-pathogenic” T_H_17 signature, as early as 3h after SAA treatment (Lee et al., 2012) (Figure 2B). Examples of increased pathogenic-T_H_17 signature genes in differentiating T_H_17-SAA1 cells include *Csf2, Tbx21 and Gzmb*, while decreased non-pathogenic-T_H_17 signature genes include *Maf, Ahr and Il10* (Figures S2D and S2E) (Lee et al., 2012). We validated elevated protein expression of GM-CSF (*Csf2*) and T-bet (*Tbx2*1) in T_H_17-SAA1-compared to T_H_17-TGFβ−differentiated cells (Figures S2F-H). SAA1 had no measurable effect in T_H_1 differentiation culture conditions (Figure S2I and S2J). Using CD4^+^ T cells from *IL23r^eGFP^* reporter mice, we confirmed that SAA1, compared to TGF-β, potently augmented IL-6-mediated *IL23r* induction (Figure 2C). This translated into stronger STAT3 activation (pSTAT3) in SAA-differentiated T_H_17 cells upon exposure to IL-23 (Figure 2D). These *in vitro* studies thus suggest that SAA1 sensitizes T_H_17 cells to inflammatory cytokines and poises them for pathological responses.

**Figure 2.**
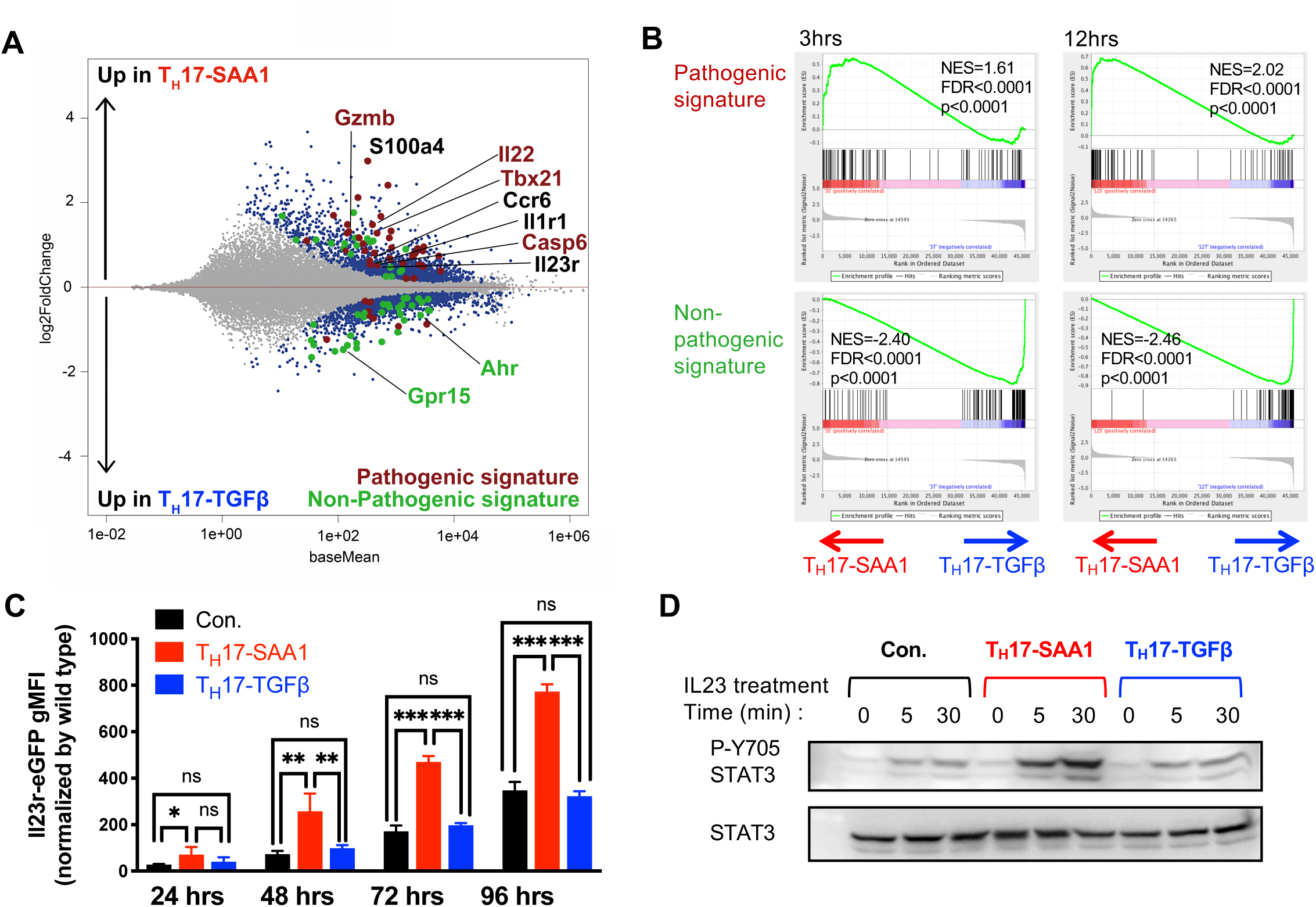
SAA1 elicits a pathogenic T_H_17 program *in vitro*. **(A and B)** RNA sequencing analysis of temporal gene expression in IL6 + αTGF-β + SAA1 (T_H_17-SAA1, n = 3) and IL6 + TGF-β (T_H_17-TGFβ, n = 3) differentiated CD4^+^ T cells. **(A)** MA plot depicts differentially expressed (DE) genes of T_H_17-SAA1 versus T_H_17-TGFβ cells at 48h *in vitro* polarization. Colored dots are significant DE genes. Red dots or green dots highlight pathogenic or non-pathogenic T_H_17 signature genes, respectively. DE genes were calculated in DESeq2 using the Wald test with Benjamini-Hochberg correction to determine the false discovery rate (FDR < 0.01). **(B)** GSEA plots of pathogenic (top) or non-pathogenic (bottom) T_H_17 signatures amongst T_H_17-SAA1 and T_H_17-TGFβ gene sets. NES, normalized enrichment score. **(C)** Summary of eGFP expression in T cells from *Il23r^eGFP^* mice following indicated condition and time of polarization. The gMFI of non-reporter, wild-type cells was subtracted from gMFI of eGFP to obtain the normalized gMFI. Representative data of two independent experiments. Statistics were calculated using the two-stage step-up method of Benjamini, Krieger and Yekutieliun. Error bars denote the mean ± s.d. ns = not significant, *p < 0.05, **p < 0.01, and ****p < 0.0001. **(D)** Amount of Y705 phosphorylated STAT3 following IL-23 treatment (10ng/ml) of T_H_17 cells differentiated under different conditions for indicated times. Total STAT3 is shown as a loading control. See also Figure **S2**.

### SAAs direct human Th17 cell differentiation and expression is associated with inflamed colon of IBD patients

Multiple genes involved in the T_H_17 pathway have been described as contributing to inflammatory bowel disease (IBD) in humans. In particular, mutations in *IL23R* result in either protection or predisposition to disease (Abdollahi et al., 2016; Duerr et al., 2006; Gaffen et al., 2014). To determine if the SAAs also contribute to human T_H_17 cell differentiation, we subjected naïve CD4^+^ T cells from cord blood to T_H_17 differentiation conditions with and without inclusion of recombinant human SAA1 or SAA2. Both SAAs substantially enhanced RORγT expression and IL-17A production in cells differentiated from multiple donors (Figures 3A-C and S3A). As in mouse, SAAs also promoted human T_H_17 cell differentiation in absence of TGF-β, based on upregulated expression of the RORγt-dependent chemokine receptor CCR6 (Figure S3B and S3C). Increased *SAA1/2* gene expression in inflamed tissues and elevated SAA serum concentrations have been reported in IBD patients with UC and CD (Lloyd-Price et al., 2019; Tang et al., 2017; Yarur et al., 2017). Using an antibody recognizing SAA1 and SAA2, we observed that SAA was prominently expressed by epithelia and lamina propria cells of inflamed but not non-inflamed adjacent regions of biopsies from UC patients (Figure 3D, 3E and S3D). Further, in a single cell RNAseq dataset of the terminal ileum lamina propria (Martin et al., 2018), there was selective heterogeneous expression of *SAA1* and *SAA2* in activated, *podoplanin*^+^ (*PDPN*) fibroblast-like cells isolated from inflamed lesions of CD patients (Figure 3F). Of note, this population was part of a highly pathogenic module characterized by an inflammatory mononuclear phagocyte (MNP)-associated cellular response organized around Ig<u>G</u> plasma cells, inflammatory <u>M</u>NP, <u>a</u>ctivated <u>T</u> and <u>s</u>tromal cells (GIMATS) that could predict resistance to anti-TNFα therapy (Martin et al., 2018) and is reminiscent of the oncostatin M receptor-expressing PDPN^+^ population identified by Powrie and colleagues that also correlated with IBD severity and resistance to anti-TNFα therapy (West et al., 2017).

**Figure 3.**
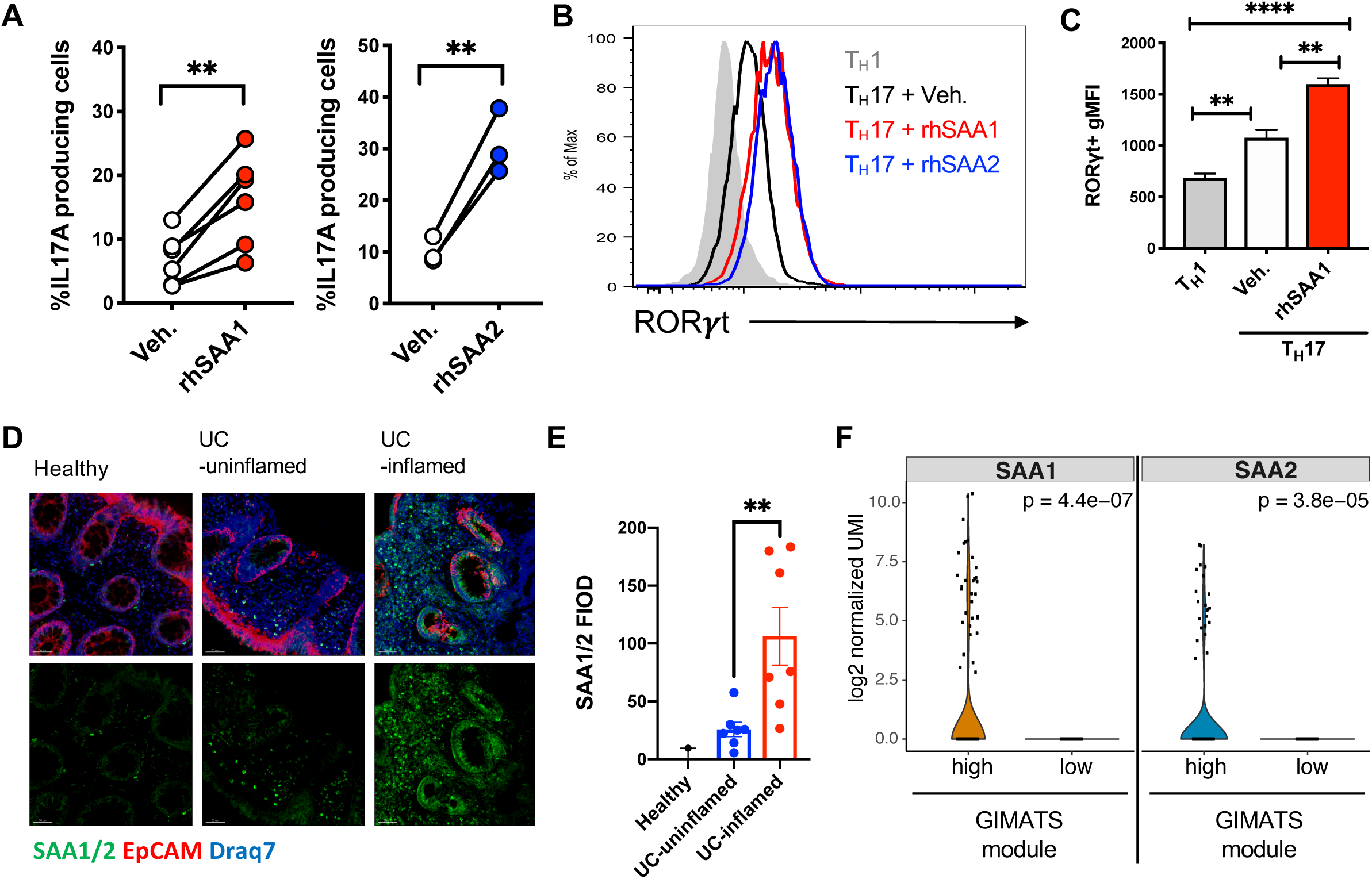
Human SAA expression in inflamed tissue and induction of T_H_17 cell differentiation. **(A-C)** Naïve human CD4^+^ T cells were isolated from cord blood and differentiated for 6 days in T_H_17 polarizing conditions ± recombinant human (rh) SAA1 or rhSAA2. **(A)** Summary of IL-17A production from in vitro polarized human T_H_17 cells. Connecting lines signify cells from the same donor. **(B)** Stacked histogram illustrates representative RORγT expression using the indicated polarizing conditions. **(C)** Summary of RORγT gMFI. Summary of 2 experiments with n = 6 donors for rhSAA1 and with n = 3 donors for rhSAA2. **(D and E)** Representative confocal images (D) and quantification of SAA1/2 by fluorescence integrated optical density (FIOD) levels (E) show prominent SAA expression in biopsies of inflamed tissue from ulcerative colitis (UC, n = 7) patients. Panels from left to right: Healthy (n = 2), UC-uninflamed, and UC-inflamed. Scale bar corresponds to 50µm. SAA1/2 (green), EPCAM (red) and nucleus (Draq7; blue). **(F)** Violin plots showing the log2 normalized UMI of SAA1 and SAA2 genes in the fibroblast cluster associated with the GIMATS module (IgG plasma cells, inflammatory MNP, activated T and stromal cells). Statistics were calculated using the paired two-tailed Student’s t-test. **p < 0.01, ****p < 0.0001. See also Figure **S3**.

### SAAs are required for differentiation and pathogenicity of colitogenic T_H_17 cells

The shared ability of mouse and human SAAs to propel *in vitro* T_H_17 cell differentiation, and the observation that SAA production was elevated in inflamed lesions in IBD patients, compelled us to examine whether SAAs exert important functions in mouse colitis models. We generated *Saa* isotype 1, 2, 3 triple-deficient (SAA^TKO^) mice and colonized these and WT littermates with *Helicobacter hepaticus* (*H. hepaticus*) (Xu et al., 2018). We then transferred into these mice naïve *H. hepaticus*-specific TCR transgenic (HH7-2Tg) CD4^+^ T cells and tracked the anti-bacterial T cell response (Xu et al., 2018) (Figure 4A and Figure S4A). Previously, we showed that T cells primed by *H. hepaticus* in WT animals differentiate into predominantly RORγt^+^Foxp3^+^ induced regulatory T cells (iTreg) or Bcl6^+^ follicular helper cells (T_FH_) (Xu et al., 2018). In this regard, examination of HH7-2Tg cells in the colonic lamina propria 2 weeks post-adoptive transfer revealed that *H. hepaticus* elicited similar numbers and frequencies of HH7-2Tg iTreg and T_FH_ cells in healthy SAA^TKO^ and WT littermate animals, indicating that SAAs did not affect T cell differentiation at steady state (Figures S4B-C). Using the same scheme, we subjected animals to continuous IL-10RA blockade, which subverts the function and maintenance of iTreg cells, resulting in expansion of cognate pathogenic T_H_17 cells and colitis (Kullberg et al., 2006; Xu et al., 2018) (Figure 4A). Sustained IL-10RA blockade resulted in abundant SAA1/2 in the serum of WT mice and upregulation of *Saa1-3* transcripts in the proximal colon, specifically, *Saa1* and *Saa2* in epithelium and *Saa3* in monocytes/macrophages and dendritic cells (Figures 4B-D and Figures S4D and S4E). By contrast, in the terminal ileum of the same mice, which were also colonized with SFB, only transcripts for *Saa1 and Saa2* were detected and there was little change upon IL-10RA blockade (Figure 4D and Figure S4F).

**Figure 4.**
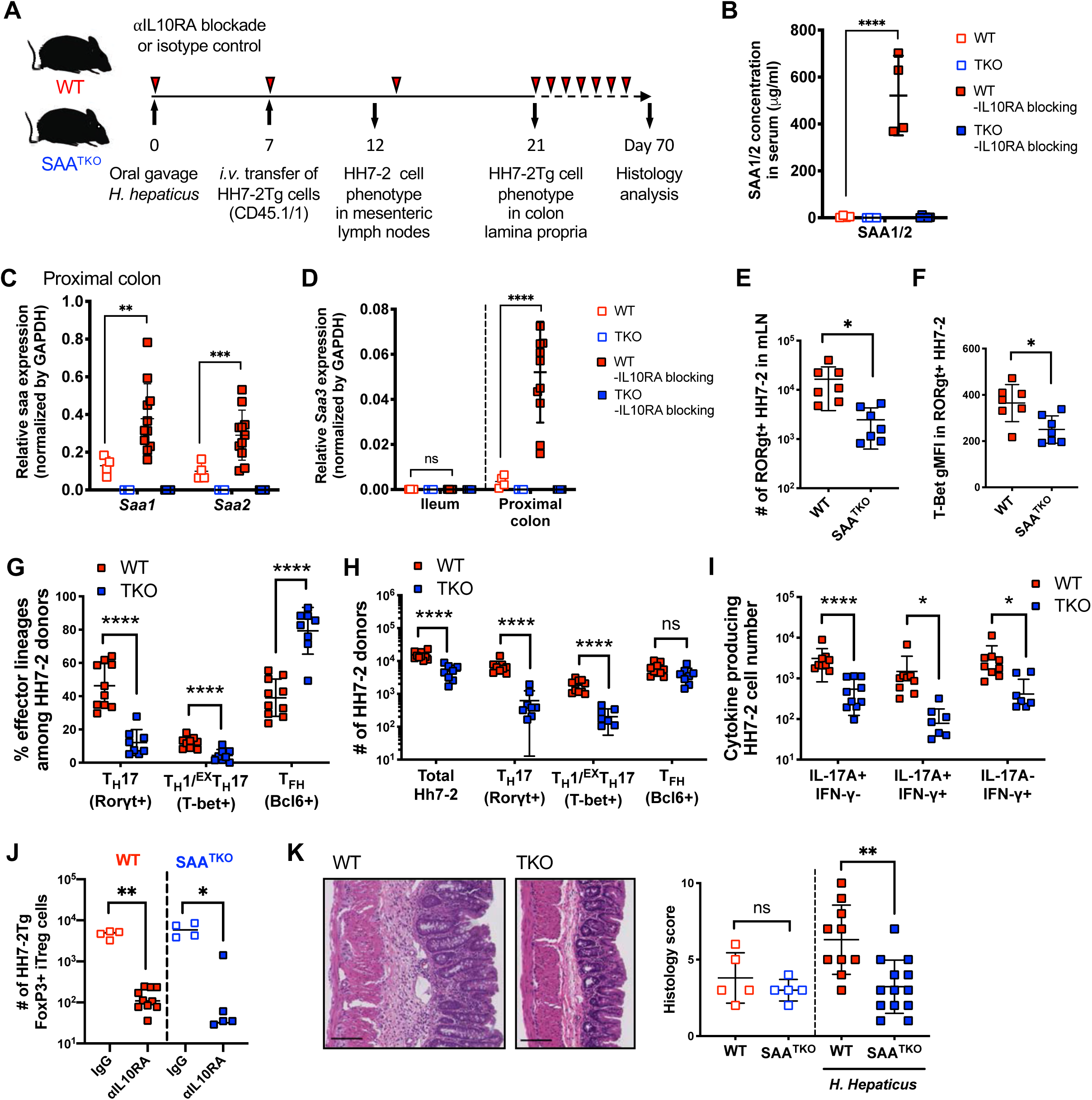
SAAs drive pathogenic T_H_ cell responses in IL-10 deficiency-dependent colitis. **(A)** Experimental scheme to examine HH7-2tg cells in SAA1/2/3 triple knock-out (SAA^TKO^) and WT littermate recipient mice colonized with *H. hepaticus* ± IL-10RA blockade (αIL10RA). **(B)** Serum concentrations of SAA1/2 of recipient mice at day 21 post *H. hepaticus* colonization. **(C and D)** Normalized expression of *Saa1/2* in proximal colon (**C**) and *Saa3* in ileum and colon (**D**) of recipient mice at day 21 post *H. hepaticus* colonization. **(E and F)** Characterization of HH7-2tg donor-derived cells 5 days post-adoptive transfer in mesenteric lymph nodes of recipient SAA^TKO^ (blue boxes, n = 8) and WT (red boxes, n = 10) littermates injected with αIL10RA. Number of RORγt expressing HH7-2 cells **(E)** and T-bet gMFI level in the RORγt^+^ HH7-2 cells **(F)**. **(G-J)** Characterization of HH7-2tg donor-derived cells two weeks post-adoptive transfer in colon lamina propria of recipient mice injected with αIL10RA. Summary of 2 experiments with SAA^TKO^ (blue boxes, n = 8) and WT (red boxes, n = 10) littermates. Frequency (**G**) and number of the indicated T_H_ cells based on transcription factor expression **(H)** or cytokine expression after restimulation (**I**). Numbers of Foxp3^+^ iTreg cells in isotype-treated recipients (hollow boxes) versus αIL10RA-treated recipients (**J**). **(K)** Representative H&E staining (left) and summary of histology scores (right) of colon sections harvested from mice with or without *H. hepaticus* colonization, following twelve weeks of αIL10RA injection. Summary of two separate experiments is shown. Uncolonized mice (open boxes): WT + αIL10RA (red, n = 5), SAA^TKO^ + αIL10RA (blue, n = 5); or colonized with *H. hepaticus* (closed boxes): WT + αIL10RA (red, n = 10), SAA^TKO^ + αIL10RA (blue, n = 13). **(B-F and K)** Statistics were calculated using the unpaired two-sided Welch’s t-test. Error bars denote the mean ± s.d. ns = not significant, **p < 0.01, ***p < 0.001, and ****p < 0.0001. **(G-J)** Statistics were calculated using the two-stage step-up method of Benjamini, Krieger and Yekutieliun. Error bars denote the mean ± s.d. ns = not significant, *p < 0.05, ***p < 0.001, and ****p < 0.0001. See also Figure **S4**.

We then assessed the HH7-2Tg cell phenotype in the mesenteric lymph nodes (mLN) of WT and SAA^TKO^ recipients at 5d after transfer. We observed reduced numbers of RORγt^+^ T_H_17 cells in SAA^TKO^ mice compared to WT littermates, in which T-bet upregulation was also markedly decreased (Figure 4E and 4F). Phenotypic analysis of HH7-2Tg cells in the colon lamina propria at two weeks after transfer revealed striking reductions in both proportion and absolute number of HH7-2Tg T_H_1 and T_H_17 cells, corresponding to profoundly decreased IFN-γ^+^, IL-17A^+^, and IFN-γ^+^IL-17A^+^ cells in SAA^TKO^ recipients (Figures 4G-I). Notably, previous IL-17A fate mapping experiments performed in mice treated with anti-IL-10RA during *H. hepaticus* infection (Morrison et al., 2013), and other pathogenic T_H_17 responses, such as experimental autoimmune encephalomyelitis (EAE), demonstrated that a significant proportion of IFNγ^+^ cells previously expressed or descended from cells that expressed the T_H_17 program (Harbour et al., 2015; Hirota et al., 2011). Therefore, reduced numbers of HH7-2Tg IFN-γ^+^ cells in SAA^TKO^ mice could be explained, at least in part, by the effect of SAAs on T_H_17 cells. Despite lower T_H_1 and T_H_17 responses in SAA^TKO^ mice, we observed a similar reduction in both the number and frequency of HH7-2Tg T_reg_ cells in both mutant and WT recipients after αIL10RA blockade (Figure 4J and Figure S4G). In accord with their reduced pathogenic T_H_ response, SAA^TKO^ mice exhibited significantly attenuated histological features of colitis compared to WT littermates following chronic IL-10RA blockade (Figure 4K). Altogether, these findings argue that the SAAs have a major role in the induction of pathogenic T_H_17 responses both *in vitro* and *in vivo*, and thus contribute to exacerbated colonic inflammation.

### Distinct sources of SAA promote experimental autoimmune encephalomyelitis at discrete stages of disease

To address the impact of SAAs on another T_H_17-dependent autoimmune disease model, we immunized mice with myelin oligodendrocyte glycoprotein (MOG) to induce EAE. SAA^TKO^ mice displayed delayed EAE onset and significantly milder disease compared to WT littermates, and had reduced T_H_17 cell accumulation in the central nervous system (CNS) (Figures 5A-C). Following MOG immunization, transcription of *Saa1* and *Saa2,* but not *Saa3*, increased in the liver, which corresponded to high concentrations of SAA1/2 in the serum beginning at the preclinical stage of disease (Figures 5D and S5A). At the peak of disease, we also detected SAA1/2 in efferent lymph, though not in the CNS (Figure S5B-C). In contrast, *Saa3* was the only *Saa* induced in the CNS during EAE, and was expressed by both microglia and monocyte-derived macrophages, particularly at the peak of disease (Figures 5E and S5D-F). Immunofluorescence staining also revealed colocalization of SAA3 with IBA1, a specific marker of microglia and monocyte-derived macrophages (Ajami et al., 2011) (Figures 5F and S5G). Thus, SAAs are broadly induced during EAE; however, their temporal and spatial sequestration suggests that they differentially influence the fate of pathogenic T_H_17 responses.

**Figure 5.**
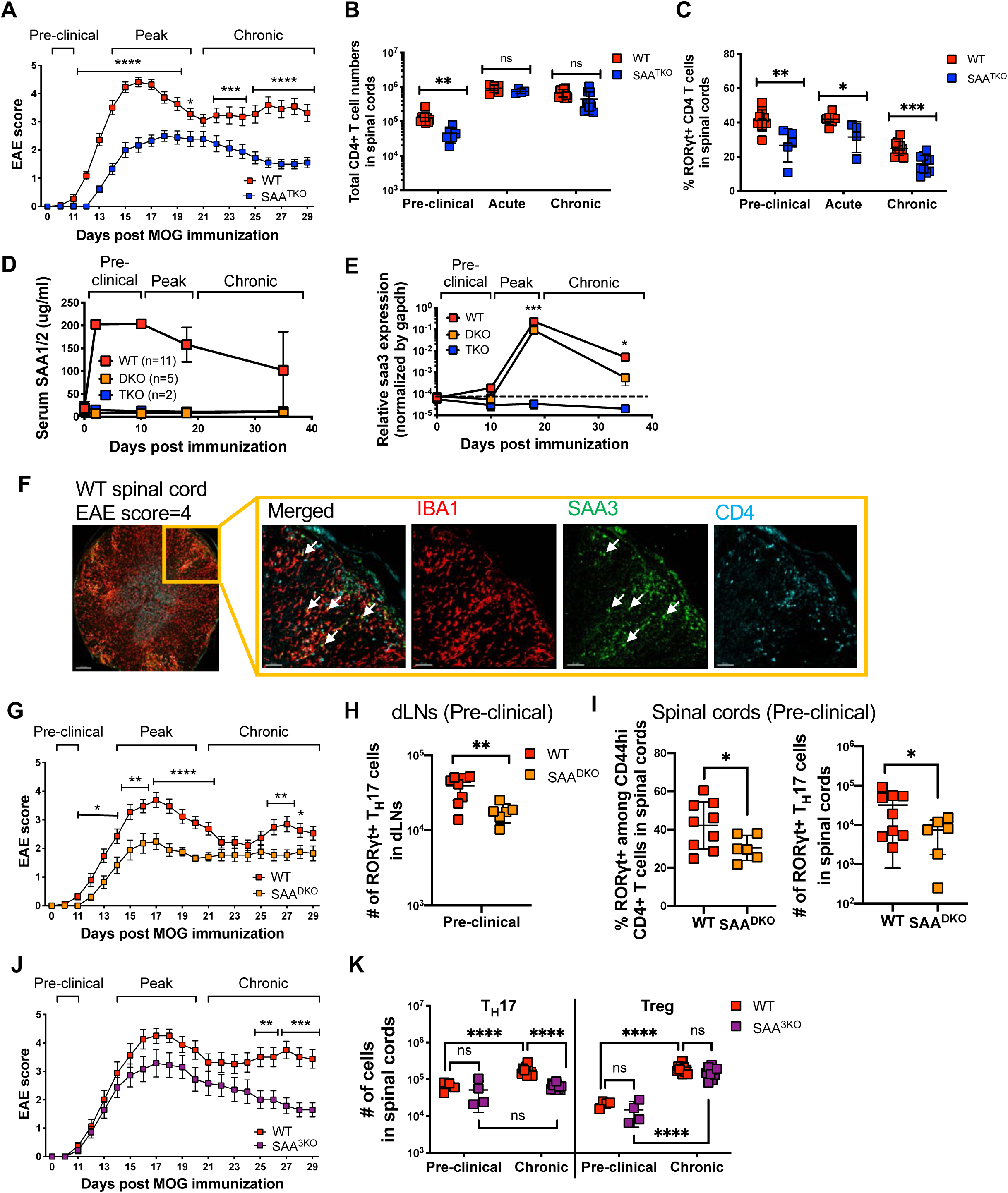
Distinct sources of SAAs promote and sustain autoimmune encephalomyelitis. **(A)** Mean EAE scores of myelin oligodendrocyte glycoprotein (MOG)-immunized SAA^TKO^ (blue boxes, n = 18) and WT littermate mice (red boxes, n = 22). Summary of 3 experiments. **(B and C)** Number of CD4^+^ T cells **(B)** and frequency of RORγt^+^ T_H_17 cells among Foxp3^Neg^CD44^hi^ CD4^+^ T cells **(C)** in the CNS of mice at the indicated stage of EAE. day 10 = pre-clinical, day 15 = acute, day 32 = chronic. Pre-clinical, (SAA^TKO^ = 6, WT = 9), acute, (SAA^TKO^ = 4, WT = 6) and Chronic, (SAA^TKO^ = 10, WT = 11). Data combined two, two, and three experiments for the pre-clinical, acute, and chronic stages of disease, respectively. **(D)** Longitudinal mean serum concentrations of SAA1/2 in MOG-immunized mice. Error bars denote the s.d. **(E)** Normalized relative expression of *Saa3* in CNS of MOG-immunized mice, measured by qPCR. Spinal cords were isolated at days 0, 10, 18, and 38 post-immunization. For WT, SAA^DKO^, and SAA^TKO^, the number of samples at each time point were 13, 7, and 13 (day 0); 5, 6, and 8 (day 10); 17, 11, and 10 (day 18); and 10, 3, 6 (day 38). **(F)** Representative confocal image of spinal cord cross section isolated from a WT mouse (n = 2) at the peak of EAE (score 4). IBA1 (red), SAA3 (green), and CD4 (aqua). White arrows pointed at yellow regions indicate IBA1/SAA3 colocalization. **(G)** Mean EAE scores of MOG-immunized SAA1/2 DKO (SAA^DKO^, orange boxes, n = 17) and WT littermate mice (red boxes, n = 19). Summary of 4 experiments. **(H and I)** Number of RORγt^+^ T_H_17 cells among CD44^hi^ effector/memory CD4^+^ T cells isolated from draining lymph nodes (dLNs) **(H)** and spinal cord **(I)** of SAA^DKO^ (n = 6) or WT (n = 9) littermates at day 10 post immunization. **(J)** Mean EAE scores of MOG-immunized SAA3 KO (SAA^3KO^, purple boxes, n = 14) and WT littermate mice (red boxes, n = 16). Summary of 3 experiments. **(K)** Number of RORγt^+^ T_H_17 and Foxp3^+^ Treg cells at pre-clinical (day 10) and chronic (day 32) stages of EAE. Summary of 3 experiments, with SAA^3KO^ (n = 14) and WT (n = 16) littermates. **(A-C, E, G, J, and K)** Statistics were calculated using the two-stage step-up method of Benjamini, Krieger and Yekutieliun. Error bars denote the mean ± s.e.m **(A, G, and J)** or mean ± s.d. **(B, C, E, H, I, and K)**. *p < 0.05, **p < 0.01, ***p < 0.001, and ****p < 0.0001. **(H and I)** Statistics were calculated using the unpaired two-sided Welch’s t-test. Error bars denote the mean ± s.d. ns = not significant, *p < 0.05, **p < 0.01, ***p < 0.001, ****p < 0.0001. See also Figure **S5**.

To distinguish the role of liver-from CNS-derived SAAs, we compared the course of EAE in SAA1 and SAA2 double-deficient (SAA^DKO^) and SAA3-deficient (SAA^3KO^) mice, respectively. Like SAA^TKO^ mice, SAA^DKO^ mice exhibited delayed disease onset and milder symptoms than WT littermates (Figure 5G). SAA^3KO^ mice, on the other hand, did not exhibit delayed disease onset; however, the magnitude of disease, particularly during the chronic stage of EAE, tapered faster in these animals than in WT littermates (Figure 5J). T_H_17 responses were also in striking concordance with these phenotypes, such that T_H_17 cell differentiation was impaired in the draining lymph nodes of SAA^DKO^ mice (Figures 5H and 5I), while the T_H_17 response in the CNS of SAA^3KO^ mice was not sustained during EAE (Figure 5K). Taken together, these findings argue that systemic SAA1 and SAA2 exert functions early in EAE pathogenesis, while SAA3 functions subsequently to sustain inflammation locally in the CNS.

### Local SAA expression fuels pathogenicity of activated T_H_17 cells

To further explore the distinct functions of SAAs in EAE pathogenesis, we employed an adoptive T cell transfer model of EAE in concert with either loss- or gain-of-function of the SAAs. We differentiated T_H_17-TGFβ cells *in vitro* from naive MOG peptide-specific 2D2 TCR transgenic T cells. When supplemented with IL-23 (T_H_17-IL-23 condition), these cells readily induce EAE in recipient mice (Lee et al., 2012). Following adoptive transfer of T_H_17-IL-23 cells into WT or SAA^DKO^ mice, all recipients developed EAE with similar kinetics and severity (Figures 6A-C). Moreover, the numbers of IL-17A^+^ 2D2 cells recovered from the CNS were indistinguishable between the two sets of recipients (Figure 6D). In contrast, when T_H_17-IL-23 2D2 cells were transferred into SAA^3KO^ recipients, the majority did not develop EAE, and those that did exhibited significantly milder symptoms than WT littermates (Mean max score: WT = 5.9, SAA^3KO^ = 2.3) (Figures 6E-H). Underscoring this impaired response was a significant reduction in the number of 2D2 cells infiltrating the CNS of SAA^3KO^ mice at the peak of disease, with lower frequencies of T_H_17 cells and fewer IL-17A^+^ cells among those (Figures S6A-C). These results suggest that SAA3, produced by microglia and monocytes during EAE, engages a feed-forward loop the fuels the T_H_17 niche in inflamed tissue. Of note, although the combination of SAA3 with IL-6 had a much less striking effect than SAA1 and IL-6 on *in vitro* T_H_17 differentiation, in other T_H_17 culture settings, such as T_H_17-TGFβ or in combination with IL-6, IL-1β, and IL-23, SAA3 potentiated T_H_17 differentiation to a similar degree as SAA1. These findings indicate that SAA3 can also signal directly in T cells (Figures S6D-G).

**Figure 6.**
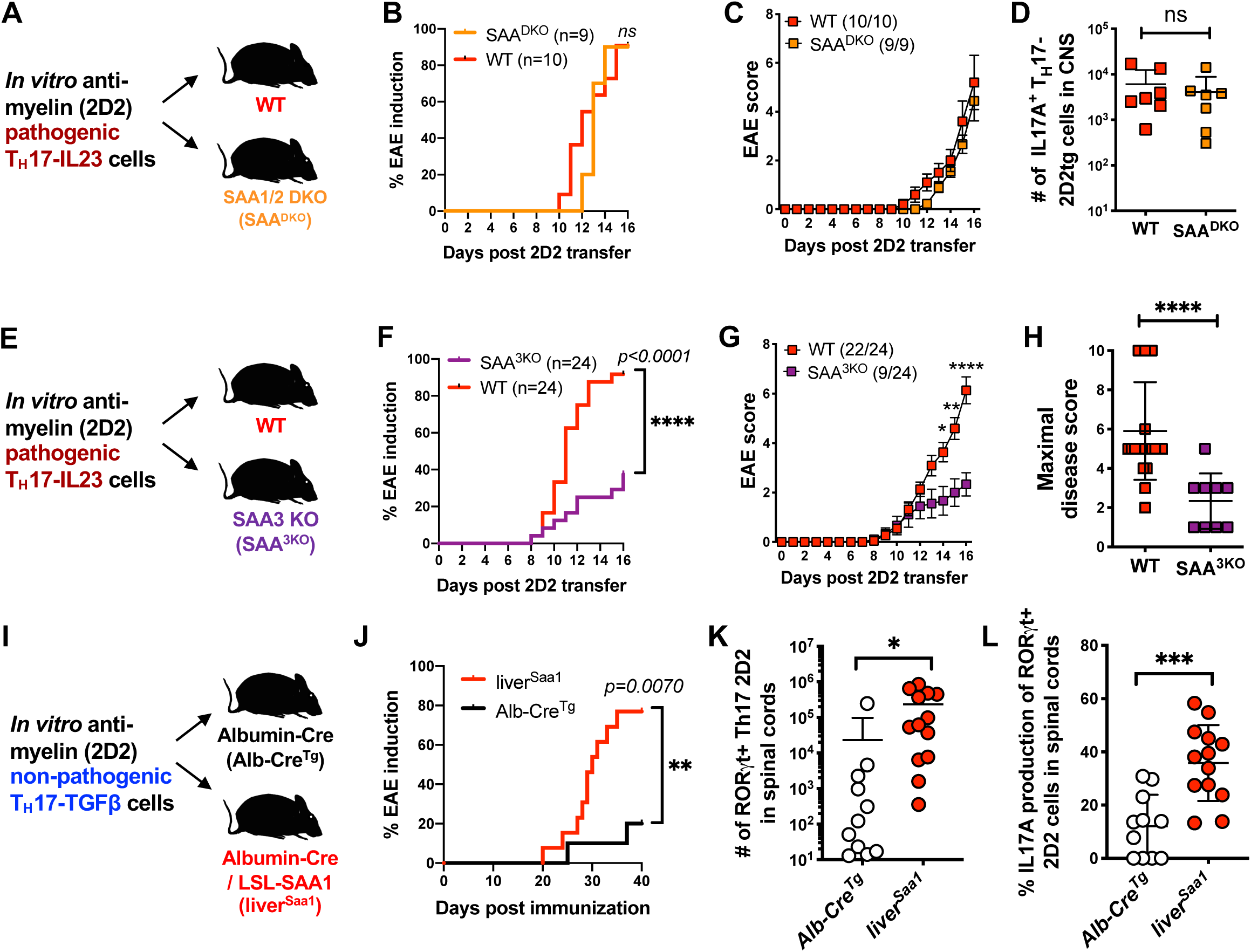
SAA regulation of pathogenicity of *in vitro*-differentiated encephalitogenic T_H_17 cells. **(A-D)** Examination of EAE development and the 2D2tg cell response in SAA^DKO^ and WT littermate recipients of 2D2tg T_H_17-IL-23 cells. Summary of 2 experiments, with SAA^DKO^ (orange, n = 9) and WT (red, n = 10) mice. Experimental scheme (**A**), EAE incidence (**B**) and mean score (**C**), and number of 2D2tg IL17A^+^ T_H_17 cells in spinal cords (**D**). **(E-H)** Examination of EAE development and the 2D2tg cell response in SAA^3KO^ and WT littermate recipients of 2D2tg T_H_17-IL-23 cells. Summary of 4 experiments, with SAA^3KO^ (purple, n = 24) and WT (red, n = 24) mice. Experimental scheme (**E**), EAE incidence (**F**) and mean score (**G**), and maximal disease score (**H**). **(I-L)** Examination of EAE development and the 2D2tg cell response in liver-specific SAA1tg (liver^SAA1^) and control Alb-Cre^tg^ littermates following transfer of 2D2tg T_H_17-TGFβ cells. Summary of 2 experiments, with liver^SAA1^ (red circles, n = 13) and Alb-Cre^tg^ (white circles, n = 11) mice. Experimental scheme (**I**), incidence of EAE onset (**J**), number of 2D2tg RORγt^+^ T_H_17 cells (**K**) and frequency of IL17A^+^ cells amongst RORγt^+^ T_H_17 cells in spinal cords at day 40 post-adoptive transfer (**L**). **(B, F, and J)** Statistics were calculated by log-rank test using the Mantal-Cox method. **(C and G)** Statistics were calculated using the two-stage step-up method of Benjamini, Krieger and Yekutieliun. Error bars denote the mean ± s.e.m. *p < 0.05, **p < 0.01, and ****p < 0.0001, **(D, H, K, and L)** Statistics were calculated using the unpaired two-sided Welch’s t-test. Error bars denote the mean ± s.d. ns = not significant, *p < 0.05, ***p < 0.001, ****p < 0.0001. See also Figure **S6**.

We next sought to probe the effect of SAAs on T_H_17 responses in the absence of other inflammatory cues. Therefore, we generated mice capable of over-expressing SAA1, by placing the coding sequence of *Saa1* downstream of a *LoxP*-flanked STOP cassette in the *Rosa26* locus (*R26^Saa1^*) (Figure S6H). When these mice expressed a *Cre* transgene under control of albumin (*Alb*-Cre^Tg^), they exhibited high concentrations of SAA1 in serum, due to constitutive production in liver. Other T_H_17 mediators, such as IL-6, were not detected (Figure S6I). Consistent with the ability of SAAs to promote T_H_17 cell differentiation *in vitro*, we found that numbers of T_H_17, but not T_H_1 cells, were increased in secondary lymphoid tissue of *Alb*-Cre^Tg^*;Rosa26^Saa1^*(liver^SAA1^) mice (Figures S6J-M). To investigate the consequence of systemically raised SAA1 levels, we transferred 2D2 T_H_17-TGFβ cells that, under normal settings, fail to efficiently evoke a pathogenic response (Figure 6I). Accordingly, few of the control *Alb*-Cre^Tg^ recipient mice developed disease (2/11), but the majority of littermate liver ^SAA1^ mice (10/13) developed EAE (Figure 6J). In addition, we recovered significantly higher numbers of RORγt^+^ T_H_17-2D2 cells, of which a greater proportion expressed IL-17A, from the CNS of liver ^SAA1^ recipients (Figures 6K and 6L). Therefore, serum SAA1/2 elevation supplants IL-23 preconditioning of T_H_17-TGFβ cells, consistent with a direct in vivo effect of SAAs on the myelin-specific T cells and suggesting that SAAs and IL-23 have convergent functions in T_H_17 autoimmune pathogenesis.

## Discussion

T_H_17 cells exert beneficial or detrimental functions under context-specific conditions at multiple body sites. In response to colonization of the small intestine with SFB, a microbe that enforces protection from enteropathogenic bacteria, T_H_17 cells promote strengthening of the epithelial barrier. Production of cytokines by SFB-specific T_H_17 cells is enhanced by local epithelial cell-derived SAA1 and SAA2, as a consequence of a signaling circuit that involves myeloid cell-derived IL-23 inducing ILC3 to produce IL-22 that, in turn, stimulates the epithelial cells. These observations prompted us to propose that T_H_17 cell effector functions are acquired in two steps, with SAA-independent priming of naïve SFB-specific T cells in the draining mesenteric lymph nodes, accompanied by their differentiation into RORγt^+^ poised T_H_17 cells, followed by migration to the lamina propria, where exposure to the SAAs results in up-regulation of IL-17 cytokines. The protective function of this type of T_H_17 cells is highlighted by the findings that blockade of IL-17A promoted epithelial barrier permeability, exacerbating DSS-induced colitis (Lee et al., 2015), and by clinical trials targeting IL-17A or IL-17RA, that failed to ameliorate CD and resulted in higher rates of adverse events or disease worsening (Hueber et al., 2012; Targan et al., 2016). In light of the previous results, which suggested that SAAs act on poised RORγt^+^ cells, we were surprised to find that SAAs act directly on naïve T cells, instructing their differentiation into T_H_17 cells in combination with the STAT3-activating cytokine IL-6. This SAA activity was critical for pathogenicity of T_H_17 cells in models of colitis and EAE, and likely reflects a program of differentiation distinct from that mediated by other combinations of cytokines both *in vitro* and *in vivo*.

### SAA-dependent differentiation of pathogenic T**H**17 cells

*In vitro*, SAA1 induced a pathogenic T_H_17 program independently of TGF-β signaling. TGF-β has been shown to be dispensable for T_H_17 differentiation under some conditions both *in vitro* and *in vivo*. For example, genetic repression of T_H_1 and T_H_2 cell differentiation was shown to relieve the requirement for TGF-β in IL-6-mediated T_H_17 induction and, indeed, T_H_17 cells can be differentiated *in vitro* by a combination of IL-6, IL-23 and IL-1β (Das et al., 2009; Ghoreschi et al., 2010). Recently, Wan and colleagues revealed that SKI, via SMAD4, transcriptionally repressed *Rorc* and that TGF-β signaling degraded or modified SKI to relieve the inhibition and promote T_H_17 cell differentiation in the presence of IL-6 (Zhang et al., 2017). Our studies indicate that SAAs function independently of STAT3 or the SMAD transcription factors, and that they do not regulate SKI degradation (unpublished). Elucidation of the receptor(s) and signaling pathways(s) engaged by the SAAs will be needed to provide insight into how they promote distinct differentiation programs for homeostatic and pathogenic T_H_17 cells.

### Context-dependent regulation of Th17 cell differentiation by SAAs

The requirement for inducible SAAs in mouse inflammatory disease models and the selective expression of SAAs in inflamed tissue from human IBD patients support a role for these secreted proteins in T_H_17-mediated inflammatory disease. Our results reveal that the effects of SAAs are subject to the context of T_H_17 cell differentiation. In SFB-mediated T_H_17 induction, the SAAs do not contribute to priming, but rather amplify the effector functions of T_H_17 cells in SFB-colonized regions. These “homeostatic” T_H_17 cells do not become pathogenic even under pro-colitogenic conditions, as occurs upon blockade of the IL-10 signaling pathway (Xu et al., 2018). By contrast, in the absence of IL-10, *H. hepaticus*-specific CD4^+^ T cells no longer adopt a T_reg_ cell fate, but instead differentiate into colitis-inducing pathogenic T_H_17 cells with a transcriptome that is strikingly different from that of homeostatic T_H_17 cells (Chai et al., 2017; Xu et al., 2018). The SAAs support this *H. hepaticus*-driven colitis, inducing pathogenic features exemplified by T-bet expression during T_H_17 priming and sensitization of the T cells to cytokines essential for T_H_17-mediated autoimmune pathogenesis, such as IL-23 and IL-1β.

The SAA proteins thus have critical roles during T_H_17 cell induction and potentiation of effector functions. It remains unclear what distinguishes these properties that are associated with pathogenic and homeostatic T_H_17 cells. It may be a feature of the microenvironment of T cell priming, e.g. in lymph nodes draining a healthy small intestine versus an inflamed large intestine, and of the other cytokines that are present; of the concentration of SAAs that are encountered; or of the cell types that produce the SAAs. A better understanding of how SAAs participate in T_H_17 cell differentiation may permit selective targeting of potentially harmful T_H_17 cells while sparing the beneficial cells.

### SAAs in mouse and human

In mouse inflammatory models, we found that SAA1 and SAA2 were prominently produced by hepatocytes after systemic immune activation and by intestinal epithelial cells adjacent to colonizing bacteria, whereas SAA3 was produced by myeloid cells in both the gut and CNS. We discovered that SAA3 advanced a feed-forward loop that sustained murine T_H_17 responses and drove EAE pathology. Although *SAA3* does not encode a full protein in humans, there are reports of inducible SAA1/2 expression by macrophages in inflamed tissues of multiple chronic inflammatory diseases (De Buck et al., 2016; Meek et al., 1994; Ye and Sun, 2015). Moreover, we and others found that tissue-derived SAA1 and SAA2 are not limited to the epithelium during inflammation. Fibroblast-like synoviocytes were shown to express and secrete SAA1/2 in rheumatoid arthritis patients (O’Hara et al., 2000), and we observed that activated fibroblasts in the inflamed parenchyma of CD patients expressed *SAA1* and *SAA2*. Given our findings and links between activated fibroblast abundance and primary anti-TNFα resistance in Crohn’s disease (West et al., 2017), it will be important to investigate whether SAAs play a causal role in this disease setting. Taken together, these findings suggest that the evolution of *SAA3* into a pseudogene in humans may be ascribed to functional redundancies with *SAA1* and *SAA2*. However, we cannot rule out that SAA3 has a unique role in the mouse, particularly since it is considerably less effective than SAA1 and SAA2 in directing T_H_17 cell differentiation in the absence of TGF-β.

### SAA signaling to T_H_17 cells in inflammatory disease

Although our results clearly demonstrate that SAAs interact directly with T cells and promote T_H_17 cell programming and response amplification, these secreted proteins also have effects on multiple other cell types. Several receptors have been described for the SAAs, including TLR2 and FPRL-1 (Ye and Sun, 2015), but inactivation of the genes for these candidate molecules had no effect on SAA1-induced Th17 cell differentiation (J-Y.L., unpublished). While we cannot rule out SAA functions through other cell types during in vivo pathogenic T_H_17 cell differentiation, our results strongly support a substantial contribution through direct interaction with T cells. In particular, the finding that enforced expression of SAA1 by hepatocytes overcame non-pathogenic T_H_17-TGFβ priming to promote EAE is consistent with sensitization of the T cells to render them more responsive to IL-23 and potentially other pro-inflammatory cytokines.

The general importance of the SAAs in maintaining the pathogenic programming of T_H_17 cells was underscored by the reductions in colitis and EAE observed when tissue SAAs were absent in mice. Elucidation of the receptor(s) for SAAs on T cells will resolve the relative contributions of direct and indirect signaling cues delivered by SAAs to T cells. However, the combined results showing that SAAs provide a previously unappreciated means for inducing inflammatory T_H_17 cells and that adoptively transferred antigen-specific T cells confer pathogenicity based on their response to SAAs argue strongly that the disease phenotypes were due, at least in part, to the breach of direct SAA interactions with T cells. Together, these findings suggest that a better understanding of T_H_17 cell responses modulated by SAAs will facilitate therapeutic efforts to ameliorate inflammatory diseases and enforce epithelial barrier integrity.

## Acknowledgements

We thank members of the Littman lab and Drs. Susan R. Schwab and Juan J. Lafaille for valuable discussions, Audrey Baeyens for collecting efferent lymph fluid from mice, Sang Y. Kim at the Rodent Genetic Engineering Core (RGEC) of NYU Medical Center (NYULMC) for generation of SAA^TKO^, SAA^3KO^ and LSL-SAA1 mice, Cindy Loomis and Experimental Pathology Research Laboratory of NYULMC for H&E staining of mouse colitis colon samples, and Adriana Heguy and the Genome Technology Center (GTC) for the RNA sequencing. The Experimental Pathology Research Laboratory is supported by National Institutes of Health Shared Instrumentation grants S10OD010584-01A1 and S10OD018338-01. The GTC is partially supported by the Cancer Center Support grant P30CA016087 at the Laura and Isaac Perlmutter Cancer Center. This work was supported by an HHMI Fellowship of the Damon Runyon Cancer Research Foundation 2232-15 (J.-Y.L.), a Dale and Betty Frey Fellowship of the Damon Runyon Cancer Research Foundation 2105-12 (J.A.H), an Alberta Innovates Fellowship (H.N.), by the Howard Hughes Medical Institute (D.R.L.), the Helen and Martin Kimmel Center for Biology and Medicine (D.R.L.), and National Institutes of Health grant R01AI121436 (D.R.L.).

## Author Contributions

J.-Y.L. and D.R.L. conceived the project; J.-Y.L. and J.A.H. performed the experiments; L.W. and H.M.S performed RNA sequencing of microglia and monocytes from EAE spinal cords; T.N. generated and purified recombinant SAA proteins; L.K. performed bioinformatic analyses; H.H.N. performed blinded histology scoring on murine colitis sections; W.-Y.L. and D.L. assisted with experiments.; S.T.Y. and K.M.K. performed tissue immunofluorescence staining; A.H., P.L., D.H., J.C.M, E.K., and M.M. provided human IBD biopsies and bioinformatics resources; J.-Y.L., J.A.H and D.R.L. wrote the manuscript.

## STAR Methods

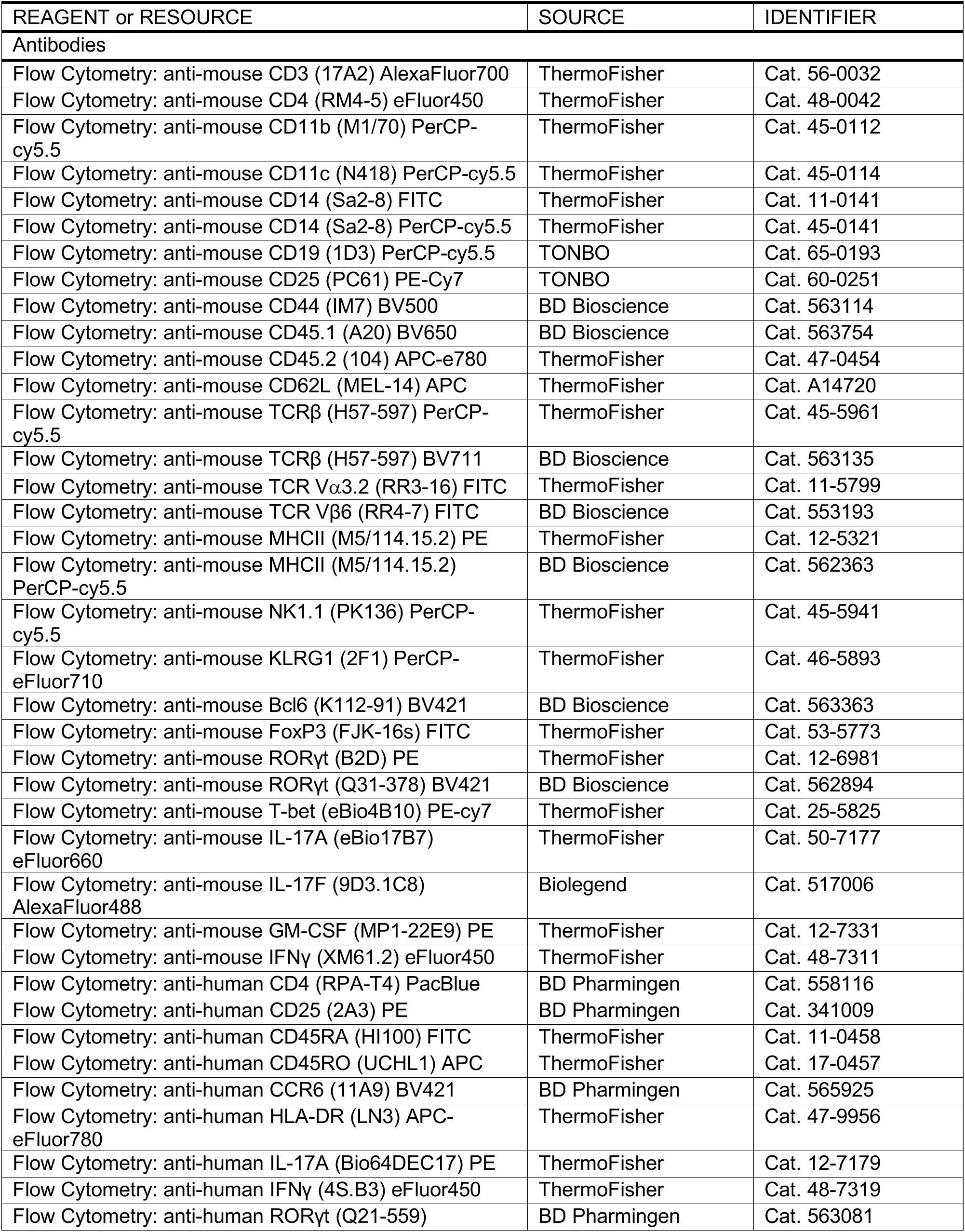
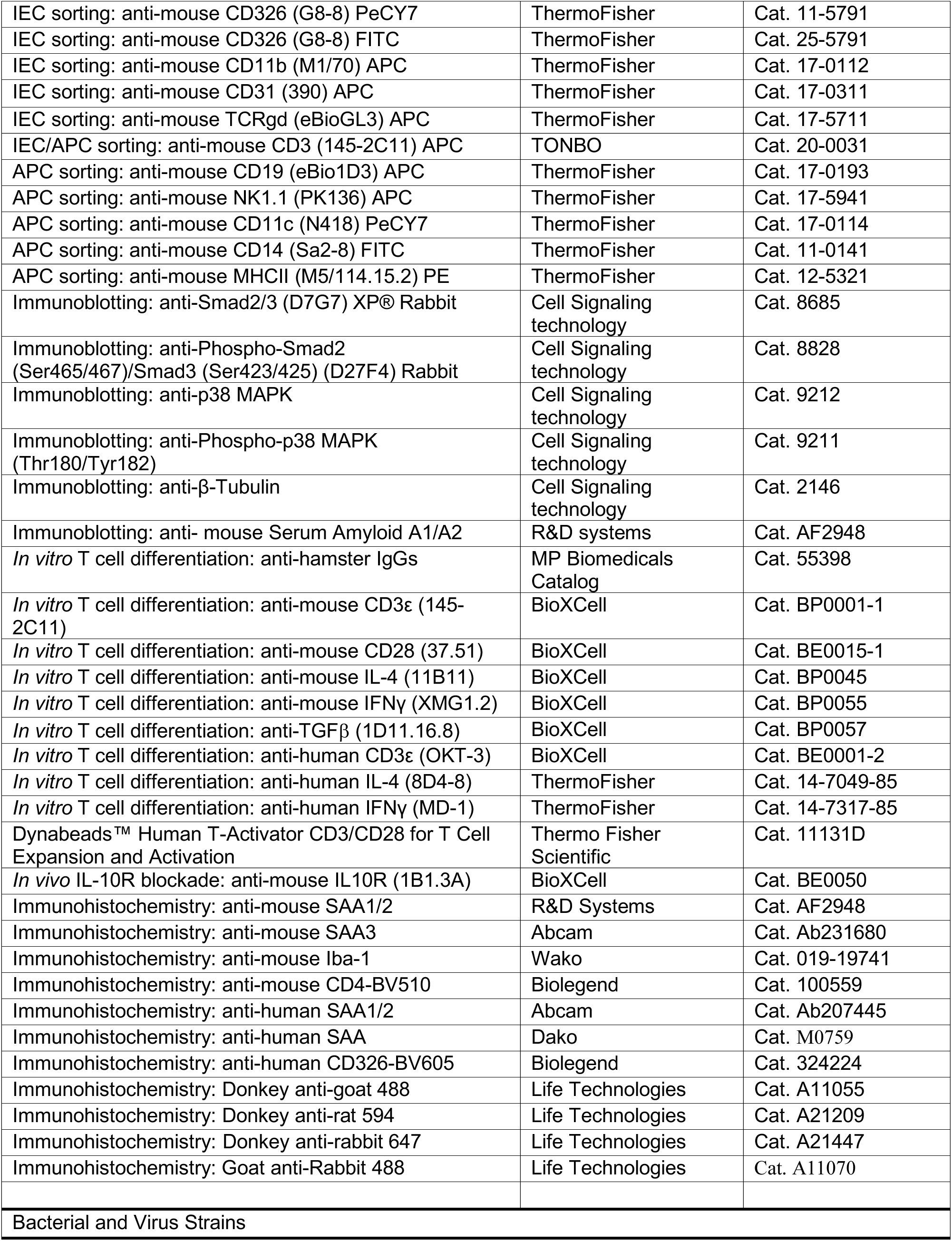
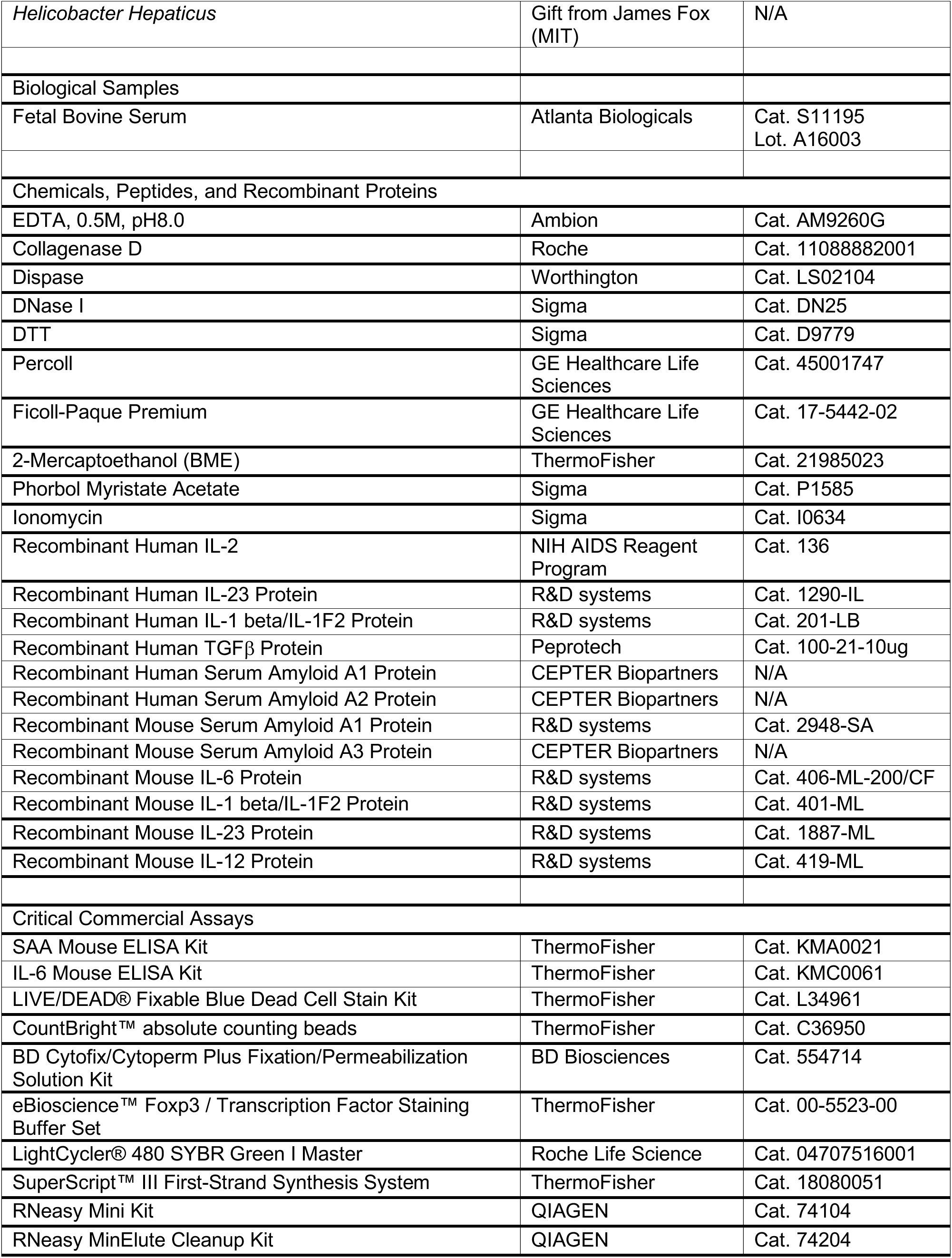
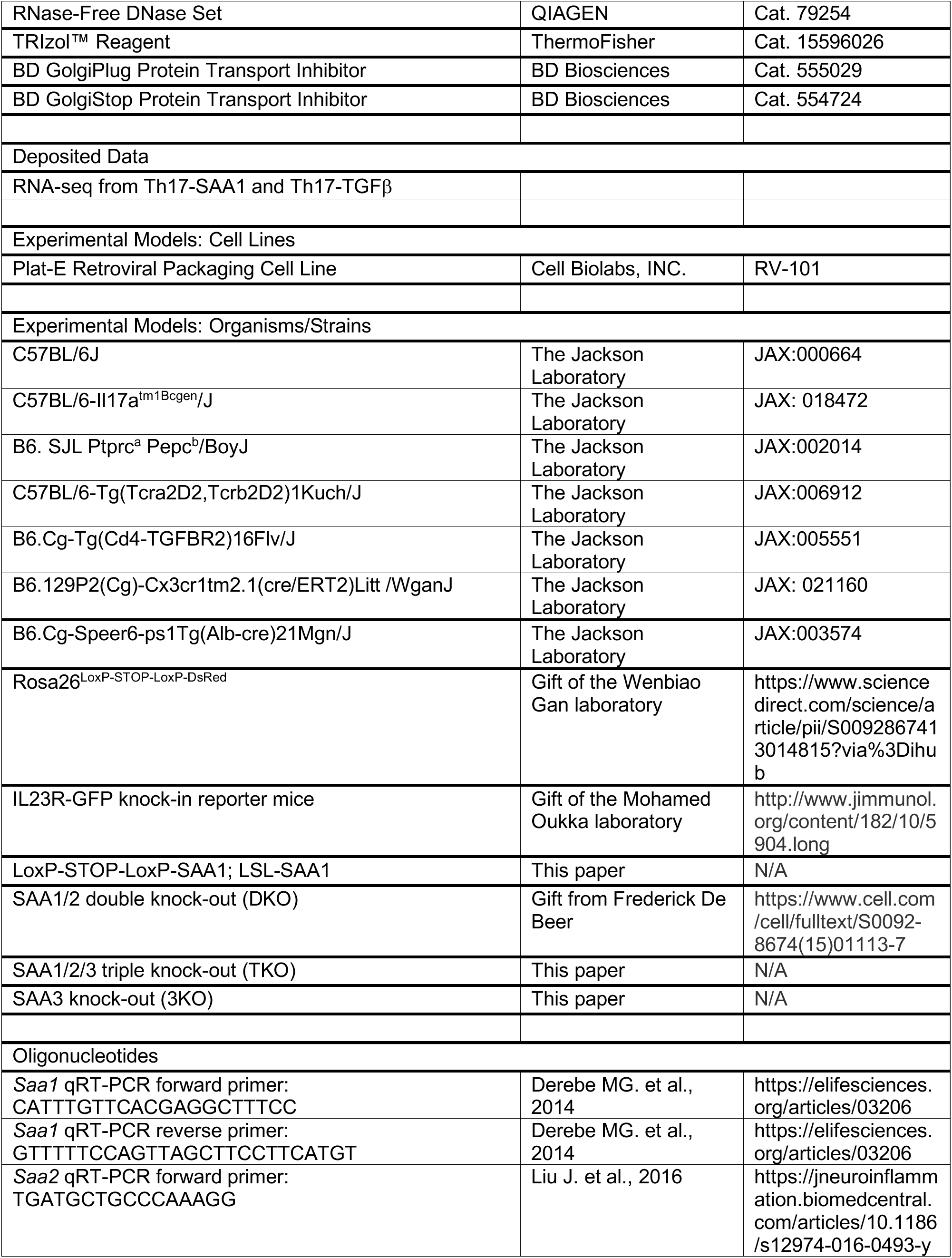
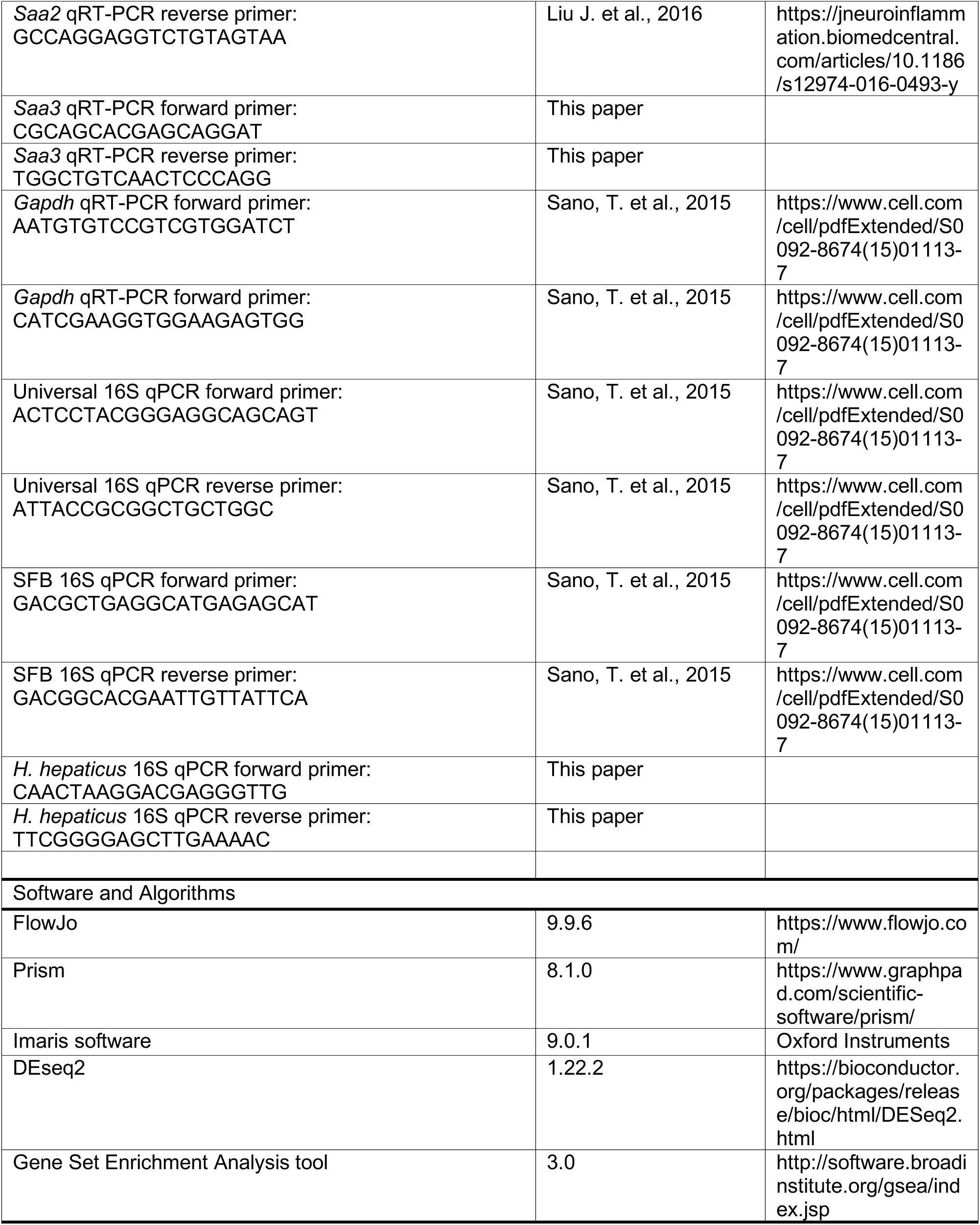
KEY RESOURCE TABLE.

### CONTACT FOR REAGENT AND RESOURCE SHARING

Further information and requests for resources and reagents should be directed to and will be fulfilled by the Lead Contact, Dan R. Littman.

### EXPERIMENTAL MODEL AND SUBJECT DETAILS

#### Mouse Strains

C57BL/6J mice were purchased from The Jackson Laboratory. All transgenic animals were bred and maintained in specific-pathogen free (SPF) conditions within the animal facility of the Skirball Institute (NYU School of Medicine. *Saa1/2* double-knockout (SAA^DKO^) mice were previously described (Eckhardt et al., 2010) and maintained bred to the Il17a-GFP reporter strain (JAX; C57BL/6-Il17a^tm1Bcgen^/J). *H.hepaticus* (Hh)-specific Th17-TCR tg (HH7) mice were previously described (Xu et al., 2018) and maintained on a Ly5.1 background (JAX; B6. SJLPtprc^a^ Pepc^b^/BoyJ). MOG-specific TCR transgenic (2D2, JAX; C57BL/6-Tg (Tcra2D2,Tcrb2D2)1 Kuch/J) mice were purchased from Jackson Laboratories, and maintained on an Ly5.1 background. CD4-dnTGFBRII (JAX; B6.Cg-Tg(Cd4-TGFBR2)16Flv/J) mice were purchased from Jackson Laboratories. Il-23r-GFP mice were provided by M. Oukka (Awasthi et al., 2009). *Cx3cr1^CreER^* (JAX; B6.129P2(Cg)-Cx3cr1^tm2.1(cre/ERT2)Litt^ /WganJ) mice were bred with Rosa26^LoxP-STOP-LoxP-DsRed^ (R26^DsRed^) mice to fate-label microglia (Parkhurst et al., 2013). Saa3 knockout (SAA^3KO^) and Saa1/2/3 triple-knockout (SAA^TKO^) mice were generated using CRISPR-Cas9 technology. A premature stop codon was inserted into exon 2 of the *Saa3* locus in WT (for SAA^3KO^) or Saa1/2 knock-out (for SAA^TKO^) zygotes. Guide RNA and HDR donor template sequences are listed in Table 1. Inducible SAA1 knock-in mice (LoxP-STOP-LoxP-SAA1; LSL-SAA1) mice were generated by targeted insertion of the STOP-eGFP-ROSA26TV cassette (Addgene Plasmid 15912: CTV vector), in which the mouse *Saa1* gene was sub-cloned, into the ROSA26 locus (Figure S6H).

**Table 1.**
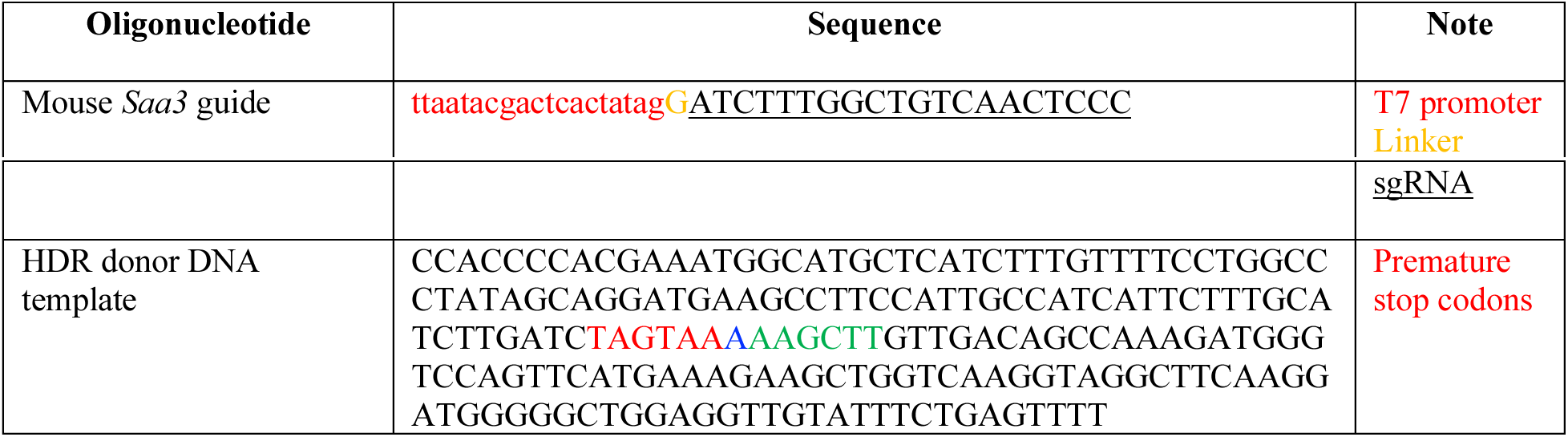
sgRNA and HDR donor sequence.

Without Cre-recombinase LSL-SAA1 mice were healthy, fertile, and born in Mendelian ratios. All in-house developed strains were generated by the Rodent Genetic Engineering Core (RGEC) at NYULMC. LSL-SAA1 mice were bred with Albumin-Cre (JAX; B6.Cg-Speer6-ps1Tg(Alb-cre)21Mgn/J) mice to generate liver-specific SAA1 overexpressing mice (LSL-SAA1/Alb-cre) (Figure S6H). Age-(6-12 weeks) and sex-(both males and females) matched littermates stably colonized with Segmented Filamentous Bacteria (SFB) were used for all experiments. To assay SFB colonization, SFB-specific 16S primers were used and universal 16S and/or host genomic DNA were quantified simultaneously to normalize SFB colonization in each sample. All animal procedures were performed in accordance with protocols approved by the Institutional Animal Care and Usage Committee of New York University School of Medicine

#### Human IBD samples

Pinch biopsies were obtained from the colon tissues (distal and rectum) of adult UC patients undergoing surveillance colonoscopy using 2.8 mm standard biopsy forceps, after protocol review and approval by the New York University School of Medicine Institutional Review Board (Mucosal Immune Profiling in Patients with Inflammatory Bowel Disease; S12-01137). All biopsies were collected in ice cold complete RPMI (RPMI 1640 medium, 10% fetal bovine serum (FBS), penicillin/streptomycin /glutamine, 50 µM 2-mercaptoethanol). Inflammation status of rectal tissue included in the study was confirmed by pathological examination as chronic active colitis.

### METHOD DETAILS

#### Flow cytometry

Single cell suspensions were pelleted and resuspended with surface-staining antibodies in HEPES Buffered HBSS. Staining was performed for 20-30min on ice. Surface-stained cells were washed and resuspended in live/dead fixable blue (ThermoFisher) for 5 minutes prior to fixation. For transcription factor staining, cells were treated using the FoxP3 staining buffer set from eBioscience according to the manufacturer’s protocol. Intracellular stains were prepared in 1X eBioscience permwash buffer containing anti-CD16/anti-CD32, normal mouse IgG (conc), and normal rat IgG (conc). Staining was performed for 30-60min on ice. For cytokine analysis, cells were initially incubated for 3h in RPMI or IMDM with 10% FBS, phorbol 12-myristate 13-acetate (PMA) (50 ng/ml; Sigma), ionomycin (500 ng/ml;Sigma) and GolgiStop (BD). After surface and live/dead staining, cells were treated using the Cytofix/Cytoperm buffer set from BD Biosciences according to the manufacturer’s protocol. Intracellular stains were prepared in BD permwash in the same manner used for transcription factor staining. Absolute numbers of isolated cells from peripheral mouse tissues in all studies were determined by comparing the ratio of cell events to bead events of CountBright™ absolute counting beads. Flow cytometric analysis was performed on an LSR II (BD Biosciences) or an Aria II (BD Biosciences) and analyzed using FlowJo software (Tree Star).

#### *In vitro* T cell culture and phenotypic analysis

Mouse T cells were purified from lymph nodes and spleens of six to eight week old mice, by sorting live (DAPI-), CD4^+^CD25^-^CD62L^+^CD44^low^ naïve T cells using a FACSAria (BD). Detailed antibody information is provided above (Flow cytometry). Cells were cultured in IMDM (Sigma) supplemented with 10% heat-inactivated FBS (Hyclone), 10U/ml penicillin-streptomycin (Invitrogen), 10μg/ml gentamicin (Gibco), 4 mM L-glutamine, and 50 µM β-mercaptoethanol. For T cell polarization, 1 x 10^5^ cells were seeded in 200 µl/well in 96-well plates that were pre-coated with a 1:20 dilution of goat anti-hamster IgG in PBS (STOCK = 1mg/ml, MP Biomedicals Catalog # 55398). Naïve T cells were primed with anti-CD3ε (0.25µg/mL) and anti-CD28 (1µg/mL) for 24 hours prior to polarization. Cells were further cultured for 48h under T_H_-lineage polarizing conditions; Control (Con. : 20 ng/mL IL-6, 2.5 µg/mL anti-IL-4, 2.5 µg/mL anti-IFNγ, 10 µg/mL anti-TGFβ), T_H_17-TGFβ (0.3 ng/mL TGF-β, 20 ng/mL IL-6, 2.5 µg/mL anti-IL-4, 2.5 µg/mL anti-IFNγ), T_H_17-SAA1 (0.15-10 µg/mL rmSAA1 (R&D systems or homemade), 20 ng/mL IL-6, 2.5 µg/mL anti-IL-4, 2.5 µg/mL anti-IFNγ, 10 µg/mL anti-TGFβ) and T_H_1 (100U/ml IL-2, 20ng/ml IL-12, 2.5 µg/mL anti-IL-4). For the SAA1 blocking assay, 2.5µg/ml of rmSAA1 was preincubated with various amount (6.25µg/ml, 12.5µg/ml, 25µg/ml, 50µg/ml) of anti-mSAA1 (R&D systems, polyclonal goat IgG, antigen affinity-purified) or control polyclonal goat IgG (R&D systems) antibodies for 30 minutes prior to the treatment. Human naïve CD4^+^ T cells were isolated from cord blood of healthy donors using anti-human CD4 MACS beads (Miltenyi), followed by CD4^+^CD25^-^HLA-DR^-^CD45RO^-^CD45RA^hi^ staining and sorting using a FACSAria (BD). Human naïve CD4^+^ T cells were cultured for 6 days in 96-well U bottom plates with 10 U/ml of IL-2, 20 ng/mL of IL-1β, 20 ng/ml of IL-23, 1 ng/ml of TGF-β, 2.5 µg/ml of anti-IL-4, 2.5 µg/mL of anti-IFNγ and anti-CD3/CD28 activation beads (LifeTechnologies) at a ratio of 1 bead per cell, as previously described (Manel et al., 2008). For monitoring cell surface expression of CCR6 on human T_H_17 cells by recombinant SAA treatment (Figure S3C and S3D), isolated human naïve CD4^+^ T cells were cultured in anti-human CD3 (OKT3)-coated 96-well flat bottom plates with 50 ng/mL of IL-1β and 50 ng/ml of IL-23, as previously described (Revu et al., 2018). 20 μg/ml of recombinant human SAA (rhSAA) proteins and 10 μg/ml anti-TGFβ neutralizing antibodies (BioXcell) were added into the culture medium during the T_H_17 differentiation.

#### Isolation of Colonic Epithelial Cells

The large intestine was removed immediately after euthanasia, carefully stripped of mesenteric fat and the cecal patch, sliced longitudinally and vigorously washed in cold HEPES buffered (25mM), divalent cation-free HBSS to remove all fecal traces. The tissue was cut into 1-inch fragments and placed in a 50ml conical containing 10ml of HEPES buffered (25mM), divalent cation-free HBSS and 2 mM of fresh DTT. The conical was placed in a bacterial shaker set to 37 °C and 200rpm for 7 minutes After 20 seconds of vigorously shaking the conical by hand, the process was repeated once more. After 20 seconds of vigorously shaking the conical by hand, the tissue was moved to a fresh conical containing 10ml of HEPES buffered (25mM), divalent cation-free HBSS and 5 mM of EDTA. The conical was placed in a bacterial shaker set to 37 °C and 200rpm for 7 minutes. After 20 seconds of vigorously shaking the conical by hand, the EDTA wash was repeated once more. Both EDTA washes were combined, passed through a 100μM filter, then spun down at 1000rpm for 5 minutes with no brake. The supernatant was carefully aspirated and the pellet resuspended in 8ml of HEPES buffered (25mM), 5% FBS-supplemented, divalent cation-free HBSS containing 4mM CaCl_2_, DNase I (100 µg/ml; Sigma), dispase (0.05 U/ml; Worthington) and then transferred into a 15ml conical. The conical was placed horizontally in a bacterial shaker set to 37 °C and 180rpm for 7 minutes. The suspension was pipetted up-down multiple times to ensure homogenization and before adding HEPES buffered (25mM), divalent cation-free HBSS to volume. The sample was spun down at 1000rpm for 5 minutes with no brake and resuspended in an Ab cocktail to enrich for IECs (See Ab table under IEC sorting) and stained for 15 minutes on ice. The cells were washed, spun down and resuspended in 2% FBS-supplemented DMEM containing DAPI and 1mM EDTA. CD326^+^ (Epcam1) cells were isolated through a 100 µM nozzle using a Sony SH800S.

#### Isolation of Lamina Propria Lymphocytes

The intestine (small and/or large) was removed immediately after euthanasia, carefully stripped of mesenteric fat and Peyer’s patches/cecal patch, sliced longitudinally and vigorously washed in cold HEPES buffered (25mM), divalent cation-free HBSS to remove all fecal traces. The tissue was cut into 1-inch fragments and placed in a 50ml conical containing 10ml of HEPES buffered (25mM), divalent cation-free HBSS and 1 mM of fresh DTT. The conical was placed in a bacterial shaker set to 37 °C and 200rpm for 10 minutes. After 45 seconds of vigorously shaking the conical by hand, the tissue was moved to a fresh conical containing 10ml of HEPES buffered (25mM), divalent cation-free HBSS and 5 mM of EDTA. The conical was placed in a bacterial shaker set to 37 °C and 200rpm for 10 minutes. After 45 seconds of vigorously shaking the conical by hand, the EDTA wash was repeated once more in order to completely remove epithelial cells. The tissue was minced and digested in 5-7ml of 10% FBS-supplemented RPMI containing collagenase (1 mg/ml collagenaseD; Roche), DNase I (100 µg/ml; Sigma), dispase (0.05 U/ml; Worthington) and subjected to constant shaking at 155rpm, 37 °C for 35 min (small intestine) or 55 min (large intestine). Digested tissue was vigorously shaken by hand for 2 min before adding 2 volumes of media and subsequently passed through a 70 µm cell strainer. The tissue was spun down and resuspended in 40% buffered percoll solution, which was then aliquoted into a 15ml conical. An equal volume of 80% buffered percoll solution was underlaid to create a sharp interface. The tube was spun at 2200rpm for 22 minutes at 22 °C to enrich for live mononuclear cells. Lamina propria (LP) lymphocytes were collected from the interface and washed once prior to staining.

For antigen presenting cell (APC) isolation, the cells were resuspended in an Ab cocktail to enrich for APCs (See Ab table under APC sorting) and stained for 15 minutes on ice. The cells were washed, spun down and resuspended in 2% FBS-supplemented DMEM containing DAPI. Dendritic cells and macrophage/inflammatory monocytes were sorted based on MHCII^+^CD11c^+^CD14^neg^ and MHCII^+^CD11c^+^CD14^+^ expression, respectively, through a 100 µM nozzle using a Sony SH800S.

For ILC3 isolation, the cells were resuspended in an Ab cocktail to remove cells positive for the following lineage markers (CD3, CD11b, CD11c, CD14, CD19, TCRβ, TCRγ, NK1.1, KLRG1). CD127^+^CD90^+^ were sorted through a 70μM nozzle on an Aria II (BD Biosciences).

#### *H. hepaticus*-induced colitis

*H. hepaticus* was provided by J. Fox (MIT) and grown on blood agar plates (TSA with 5% sheep blood, Thermo Fisher) as previously described (Xu et al., 2018). *H. hepaticus*-induced colitis was induced as described (Kullberg et al., 2006). Briefly, mice were colonized with *H. hepaticus* by oral gavage on days 0 and 4 of the experiment. 1 mg of an IL-10R-blocking antibody (clone 1B1.2) was administered by intraperitoneal injection once weekly starting at day 0. After 7 days, naïve HH7-2 TCRtg CD4^+^ T cells were adoptively transferred into the *H. hepaticus* colonized recipients. Briefly, spleens from donor HH7-2 TCRtg mice were collected and mechanically disassociated. Red blood cells were lysed using ACK lysis buffer (Lonza). Naive HH7-2 Tg CD4^+^ T cells were sorted as CD4^+^CD3^+^CD44^lo^CD62L^hi^CD25^−^Vβ6^+^ on an Aria II (BD Biosciences). Cells were resuspended in ice-cold PBS and transferred into congenic recipient mice via intravenous injection. In order to confirm *H. hepaticus* colonization, *H. hepaticus*-specific 16S primers were used on DNA extracted from fecal pellets. Universal 16S were quantified simultaneously to normalize *H. hepaticus* colonization of each sample.

To score colitis severity, colon and cecum samples were surgically removed from all groups of mice at 12 weeks post IL-10R-blockade injection. The samples were gently swiss rolled from the distal end and fixed in 4% paraformaldehyde (Electron Microscopy Science, Hatfield USA). Formalin-fixed tissues were then processed for paraffin embedding, cut into 5-micron thick sections and stained with hematoxylin and eosin (H&E) as per protocol by the Experimental Pathology Core Laboratory at New York University (NY, USA). H&E scoring was performed in a blinded fashion using coded slides. Total scores for colonic and cecum inflammation were comprised of individual scores from 4 categories: 1) Goblet cells per High Power Field (HPF) (Score of 1: 11 to 28 goblet cells per HPF; Score of 2: 1 to 10 goblet cells per HPF; Score of 3: ≤ 1 goblet cell per HPF). 2) Submucosa edema (Score of 0: no pathological changes; Score of 1: Mild edema accounting for <50% of the diameter of the entire intestinal wall; Score of 2: moderate edema involving 50-80% of the diameter of the entire intestinal wall; Score of 3: profound edema involving >80% of the diameter of the entire intestinal wall). 3) Inflammatory Infiltration Depth (Score of 0: No infiltrate; Score of 1: Infiltration above muscularis mucosae; Score of 2: Infiltration extending to include submucosa; Score of 3: Transmural Infiltration). 4) Epithelial Changes (Score of 0: No changes; Score of 1: Upper third or only surface epithelial missing; Score of 2: Moderate epithelial damage with intact base of crypts; Score of 3: Severe with missing crypts).

#### Induction of EAE by MOG-immunization or 2D2 transfer

For induction of active experimental autoimmune encephalomyelitis (EAE), mice were immunized subcutaneously on day 0 with 100µg of MOG_35-55_ peptide, emulsified in CFA (Complete Freund’s Adjuvant supplemented with 2mg/mL Mycobacterium tuberculosis H37Ra), and injected i.p. on days 0 and 2 with 200 ng pertussis toxin (Calbiochem). For 2D2 transfer EAE experiments, *in vitro* polarized 2D2 Th17 cells were injected intravenously into recipient mice at 3 × 10^6^ IL-17A producing 2D2 cells per recipient. Naive 2D2 TCR-transgenic CD4^+^ T cells from the spleen and lymph nodes of 2D2 mice were electronically sorted (CD4^+^Vβ11^+^CD62L^hi^) and activated under T_H_17-polarizing conditions, in the presence of irradiated wild-type splenocytes at a 5:1 ratio, with the following: anti-CD3 (2.5µg/ml) (145-2C11, BioXCell), anti-IL-4 (20µg/ml) (11B11, BioXCell), anti-IFN-γ (20µg/ml) (XMG1.2, BioXCell), mIL-6 (30ng/ml) (Miltenyi Biotec) and hTGF-β1 (3ng/ml) (Miltenyi Biotec). After 48 h of activation, mIL-23 (10 ng/ml; R&D Systems) was added, as necessary. On day 5 of culture, cells were reactivated on plates precoated with 2 µg/ml of anti-CD3 and anti-CD28 (PV1, BioXCell) for an additional 48 h, before adoptive transfer. The EAE scoring system was as follows: 0-no disease, 1-Partially limp tail; 2-Paralyzed tail; 3-Hind limb paresis, uncoordinated movement; 4-One hind limb paralyzed; 5-Both hind limbs paralyzed; 6-Hind limbs paralyzed, weakness in forelimbs; 7-Hind limbs paralyzed, one forelimb paralyzed; 8-Hind limbs paralyzed, both forelimbs paralyzed; 9-Moribund; 10-Death. For isolating mononuclear cells from spinal cords during EAE, spinal cords were mechanically disrupted and dissociated in RPMI containing collagenase (1 mg/ml collagenaseD; Roche), DNase I (100 µg/ml; Sigma) and 10% FBS at 37 °C for 30 min. Leukocytes were collected at the interface of a 40%/80% Percoll gradient (GE Healthcare). Efferent lymph fluids of pre-clinical stage mice were collected from cysterna chyli as previously described (Matloubian et al., 2004).

#### Microglia and monocyte-derived macrophage isolation from EAE spinal cords

*Cx3cr1^CreERT2/+^/R26^DsRed/+^* mice were gavaged with 10mg of tamoxifen diluted in corn oil at the age of 28 and 30 days. On day 60 EAE was induced by MOG-immunization and pertussis toxin injection as described. Mice were evaluated daily for weight loss and clinical development of hind limb paralysis. Mice were euthanized at the pre-clinical (day 9-10 post immunization), peak (acute; day 15-17), and chronic (day 25) stages of disease. Spinal cords were harvested, minced and digested as described. After obtaining a single cell suspension, samples were stained with the appropriate antibodies. Microglia cells and infiltrating monocyte-derived macrophage cells were sorted as DsRed^+^CX3CR1^+^CD11b^+^CD45^int^MHCII^lo^B220^-^CD3^-^ (microglia) and DsRed^-^CX3CR1^+^ CD11b^+^CD45^hi^F480^+^MHC^hi^B220^-^CD3^-^ (monocyte-derived cells) on the Aria II (BD Biosciences) respectively.

#### Tissue Preparation for Immunofluorescence, Confocal Microscopy, and Image Analysis

Tissue preparation for immunofluorescence, confocal microscopy, and image analysis was conducted as described (Perez et al., 2017). Briefly, tissues were fixed in paraformaldehyde, lysine, and sodium periodate buffer (PLP, 0.05 M phosphate buffer, 0.1M L-lysine, p.H. 7.4, 2 mg/mL NaIO4, and 10 mg/mL paraformaldehyde) overnight at 4°C followed by 30% sucrose overnight at 4°C and subsequent OCT media embedding. 20-μm frozen tissue sections were sectioned using a Leica CM3050S cryostat. FcR block was with anti-CD16/32 Fc block antibody (Biolegend) diluted in PBS containing 2% goat or donkey serum and 2% fetal bovine serum (FBS) for 1 hour at room temperature. Sections were stained with the indicated antibodies (Tables 2 and 3) diluted in PBS containing 2% goat or donkey serum, 2% FBS, and 0.05% Fc block for 1 hour at room temperature. For the intracellular staining, all antibodies including the Fc block were diluted in PBS containing 2% goat or donkey serum, 2% FBS, and 0.1% Triton-X. Images were acquired using a Zeiss LSM 880 confocal microscope (Carl Zeiss) with the Zen Black software. The imaging data were processed and analyzed using Imaris software version 9.0.1 (Bitplane; Oxford Instruments). Human SAA1/2 confocal images were analyzed by Image J software (NIH, Bethesda, MD) to measure signal intensity. Average signal intensity for each region of interest was normalized to healthy control.

**Table 2.** Antibodies and staining conditions for mouse tissues.

| Antibody | Species | Conc. | Clone | Catalog # | Lot # | Vendor |
| --- | --- | --- | --- | --- | --- | --- |
| SAA1/2 | Goat | 1:300 |  | AF2948 | VWJ0417091 | R&D Systems |
| SAA3 | Rat | 1:50 |  | Ab231680 | GR32218061 | Abcam |
| Draq-7 |  | 1:1000 |  | 564904 |  | R&D Systems |
| FC Block |  | 1:200 | 93 | 101302 | 3255480 | Biolegend |
| Donkey anti-goat 488 |  | 1:300 |  | A11055 | 1827671 | Life Technologies |
| Donkey anti-rat 594 |  | 1:300 |  | A21209 | 1979379 | Life Technologies |
| Iba-1 | Rabbit | 1:500 |  | 019-19741 | LKH4161 | Wako |
| Donkey anti-rabbit 647 |  | 1:300 |  | A21447 | 1977345 | Life Technologies |
| CD4-BV510 | Rat | 1:100 | RM4-5 | 100559 | B227810 | Biolegend |

**Table 3.** Antibodies and staining conditions for human tissues.

| Antibody | Species | Conc. | Clone | Catalog # | Lot # | Vendor |
| --- | --- | --- | --- | --- | --- | --- |
| SAA | Rabbit | 1:500 |  | Ab207445 | GR270796-2 | Abcam |
| SAA | Mouse | 1:50 | mc1 | M0759 | 20059191 | Dako |
| CD326-BV605 | Mouse | 1:100 | 9C4 | 324224 | B251369 | Biolegend |
| Draq-7 |  | 1:1000 |  | 564904 |  | R&D Systems |
| Goat anti-Rabbit 488 |  | 1:300 |  | A11070 | 1775509 | Life Technologies |
| FC Block |  | 1:200 |  | 14-9161-73 | 4350496 | Invitrogen |

#### Expression and purification of mouse recombinant SAAs

Escherichia coli codon optimized DNA sequences, encoding SAA isotypes (without the signal sequence), were chemically synthesized (GenScript) and cloned into the pET26(b)+ (Novagen) expression vector between NdeI and HindIII restriction endonuclease sites, with a C-terminal hexa-histidine. Proteins were purified following the previously published protocol (Derebe et al., eLife 2014;3:e03206) with slight modifications. Briefly, proteins were overexpressed in *E. coli* BL21(DE3) cells by induction with 0.4 mM isopropyl-β-D-galactoside (IPTG) for ∼4 hr at 25°C for mSAA1, and at 37°C for mSAA3. Cells were harvested by centrifugation at 5000×g for 20 min at 4°C and re-suspended in ice-cold lysis buffer (50 mM NaH2PO4, 500 mM NaCl, 10 mM imidizole for mSAA1 and 500 mM NaCl, 50 mM Tris pH 8.0, 10 mM imidazole, 15 mM β-mercaptoethanol for mSAA3). After sonication of cell suspension, decyl maltopyranoside (DM) (Avanti Polar Lipids) was added to a final concentration of 40 mM and incubated for ∼3 hr at 4°C followed by centrifugation at 14,000×g for 45 min. The supernatant was collected and incubated with Ni-NTA resin (GE HealthCare) pre-equilibrated with ice-cold corresponding lysis buffer containing 4 mM DM (DM buffer) for 3 h at 4 °C under mild mixing conditions to facilitate binding. The column was washed thrice with 25 mM imidazole in DM buffer to remove the non-specific contaminants. Finally, proteins were eluted using an imidazole gradient (in DM buffer) under gravity flow. Fractions containing the protein of interest were pooled and dialyzed against 20 mM Tris-HCl (pH 8.2) containing 500 mM NaCl, 0.1% (w/v) PEG 3350 and 20 mM Imidazole. The dialyzed protein was concentrated in an Amicon stirrer cell apparatus (Millipore) to a final concentration of ∼1 mg/mL. Protein purity was assessed by SDS/PAGE.

#### RNA isolation and cDNA preparation

Total RNAs from *in vitro* polarized T cells or sorted cell populations were extracted using TRIzol (Invitrogen) followed by DNase I (Qiagen) treatment and cleanup with RNeasy MinElute kit (Qiagen) following manufacturer protocols. For total tissue RNA isolation, dissected tissues were homogenized in TRIzol. cDNA was generated using a SuperScript™ III First-Strand Synthesis System (ThermoFisher).

#### qRT-PCR and qPCR

Quantitative RT-PCR and PCR were performed using the Hot Start-IT SYBR Green (Affymetrix) on the Roche real-time PCR system (Roche 480). For analysis of mRNA transcripts, RNA samples were treated with DNase (Roche) prior to cDNA synthesis to avoid effect of DNA contamination. Gene specific primers spanning exons were used. Values were normalized to GAPDH for each sample.

#### Library preparation for RNA sequencing

RNA-seq libraries for in vitro polarized T_H_17 lineages were prepared with the TruSeq Stranded Total RNA Library Prep Gold Kit (Illumina, 20020598). The sequencing was performed using the Illumina NovaSeq or NextSeq. RNA-seq libraries were prepared and sequenced by the Genome Technology Core at New York University School of Medicine. RNAseq libraries for isolated microglia and CNS-infiltrating monocytes were prepared and sequenced by the Illumina RapidRun at the Genome Services Laboratory, HudsonAlpha.

### QUANTIFICATION AND STATISTICAL ANALYSIS

#### Transcriptome analysis

Fastq files were aligned to the mouse Ensemble genome GRCm38 with STAR v 2.6.1d. Read pairs were counted using featurecounts from the Subread package v 1.6.2, prior to normalization and differential expression analysis which were performed using DESeq2. Gene set enrichment analyses were performed with the Broad Institute’s GSEA tool. Potential enrichment of pathogenic and non-pathogenic gene sets defined from previous studies (Lee et al., 2012) were analyzed with the RNAseq data at each time point.

The single cell RNA sequencing data of SAA1 and SAA2 genes in the fibroblast clusters was analyzed as described in (Martin et al., 2018).

#### Statistical analysis

Differences between groups were calculated using the unpaired two-sided Welch’s t-test or the two-stage step-up method of Benjamini, Krieger and Yekutieliun. For EAE disease induction, log-rank test using the Mantal-Cox method was performed. For RNA-seq analysis, differentially expressed genes were calculated in DESeq2 using the Wald test with Benjamini–Hochberg correction to determine the FDR. GSEA p value was calculated with gene set permutations. Genes were considered differentially expressed with FDR < 0.01 and log2 fold change > 1.2. Data was processed with GraphPad Prism, Version 8 (GraphPad Software). We treated less than 0.05 of p value as significant differences. ∗p < 0.05, ∗∗p < 0.01, ∗∗∗p < 0.001, and ∗∗∗∗p < 0.0001. Details regarding number of replicates and the definition of center/error bars can be found in figure legends.

## Supplemental Information

### Supplemental figure legends

**Figure S1.**
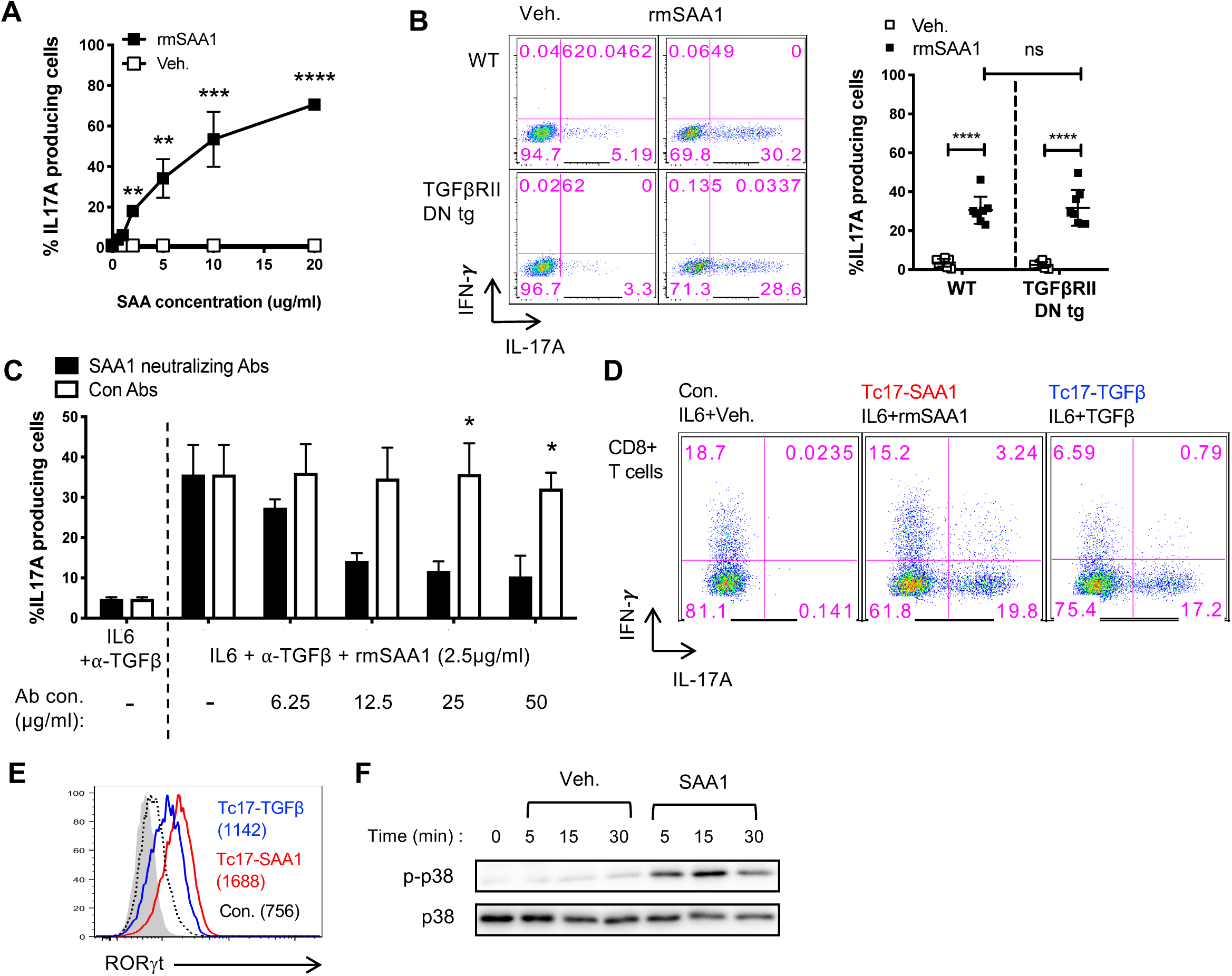
Mouse T_H_17/Tc17 cell differentiation by SAAs, Related to Figure 1. **(A)** IL-17A dose response curve to rmSAA1. **(B)** Representative IL-17A versus IFNγ expression (left) and summary (right) in WT and TGFβRII DNtg CD4^+^ T cells following SAA1-induced differentiation. **(C)** Dosage-dependent inhibition of rSAA1-induced Th17 cell differentiation by SAA1-neutralizing antibody. **(D, E)** Representative IL-17A versus IFNγ expression (**D**) and RORγt expression (**E**) in CD8^+^ T cells cultured under SAA1 or TGF-β differentiation conditions. gMFI of RORγt is in parentheses. **(F)** Immunoblotting for p38 MAPK phosphorylation (p-p38) at T180/Y182 of primed CD4^+^ T cells upon rmSAA1 (10μg/ml) or vehicle (Veh.) treatment for indicated times. Total p38 is shown as a loading control. A summary of 2 experiments with at least 2 biological replicates per experiment. **(A-E)** Experiments were conducted based using the same scheme as in Figure 1A. **(A-C)** All statistics were calculated using the unpaired two-sided Welch’s t-test. Error bars denote the mean ± s.d. ns = not significant, *p < 0.05, **p < 0.01, ***p < 0.001, ****p < 0.0001.

**Figure S2.**
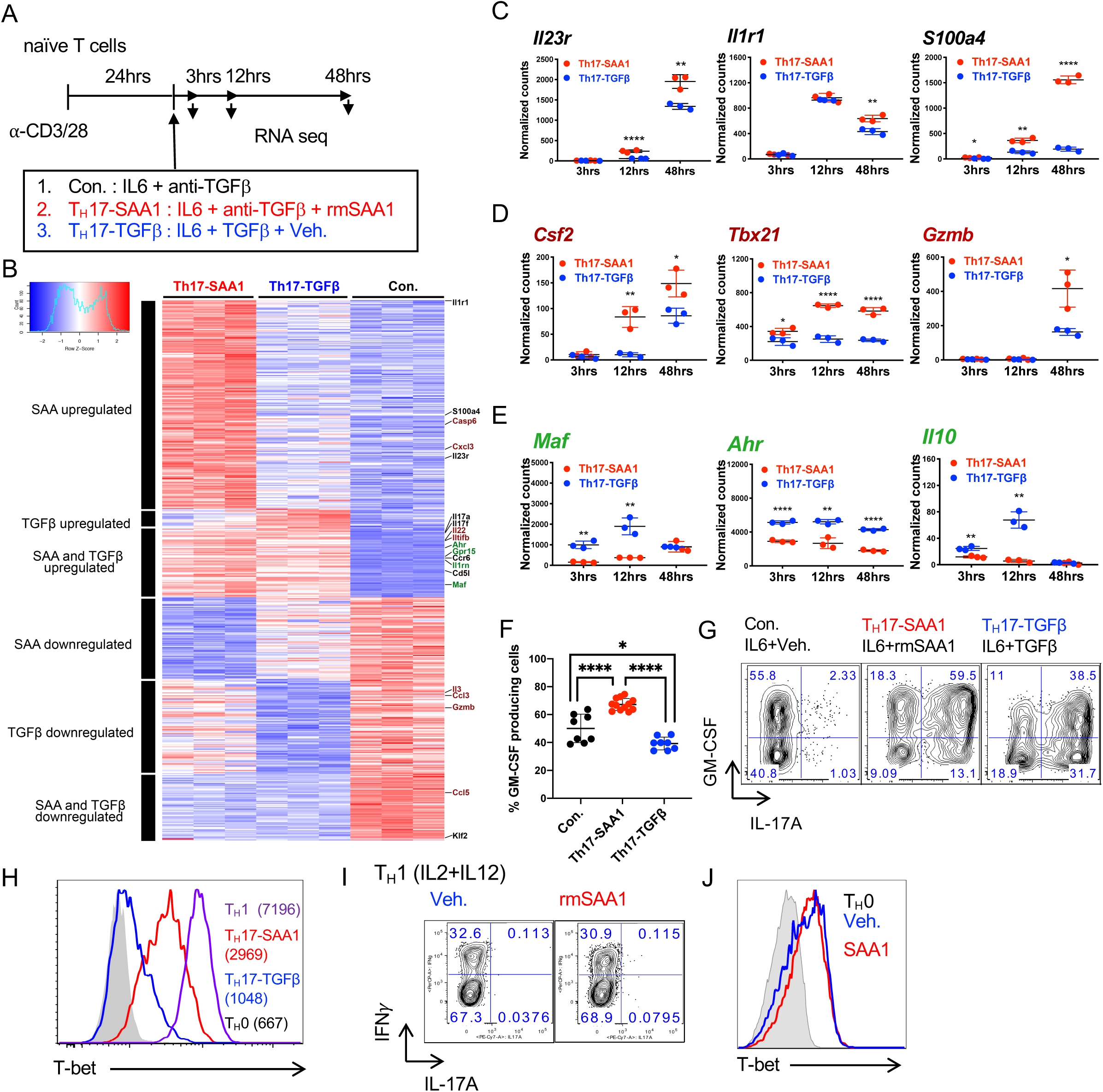
Transcriptional responses and validation of differentially expressed genes in T_H_17 cells differentiated with IL6 plus SAA1 or TGF-β, Related to Figure 2. **(A)** Experimental scheme. Cells were harvested and RNA extracted at 3, 12 and 48 h after polarization. **(B)** Clustered heatmap of differentially expressed genes at 48 h timepoint. Color scale is based on normalized read counts. Genes listed on the righthand margin are color coded. Green = non-pathogenic T_H_17 signature. Maroon = pathogenic T_H_17 signature. Black = Autoimmune disease associated. **(C-E)** Normalized counts of autoimmune disease-associated (*Il23r, Il1r1, S100a4*) **(C)**, pathogenic (*Csf2*, *Tbx21, Gzmb*) **(D)**, or non-pathogenic (*Maf, Ahr*, *Il10*) **(E)** genes at the indicated time points. **(F-H)** GM-CSF **(F and G)** and T-bet **(H)** expression at 48 h. gMFI of T-bet is in parentheses. **(I and J)** Representative expression of IFNγ versus IL-17A (I) and T-bet (J) in T_H_1 cells induced *in vitro* with IL-2 and IL-12, with and without SAA1, for 48 hrs. **(C-E)** Statistics were calculated using the two-stage step-up method of Benjamini, Krieger and Yekutieliun. Error bars denote the mean ± s.d. **(F)** Statistics were calculated using the unpaired two-sided Welch’s t-test. Error bars denote the mean ± s.d. ns = not significant, *p < 0.05, ***p < 0.001, and ****p < 0.0001.

**Figure S3.**
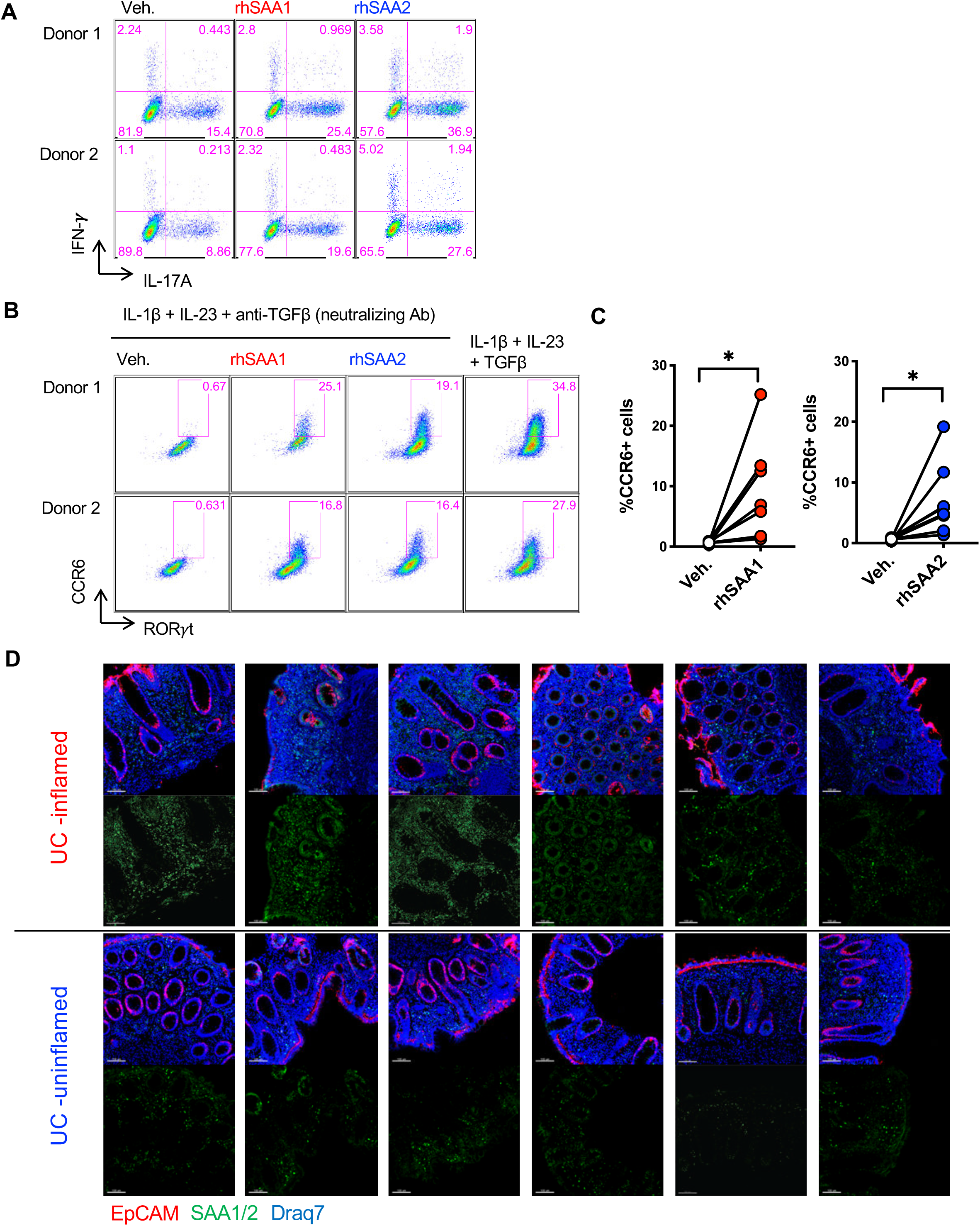
Role of SAA in human T_H_17 cell differentiation and protein expression in inflamed colon tissue from UC patients, Related to Figure 3. **(A)** Representative FACS plots of IL-17A versus IFNγ expression by *in vitro* polarized human T_H_17 cells upon restimulation. **(B and C)** Representative FACS plots (B) and summaries (C) indicating cell surface expression of CCR6 on *in vitro* polarized human T_H_17 cells upon human recombinant SAA treatment. Connecting lines signify the same donor. Summary of 3 experiments with n = 8 donors. Statistics were calculated using the paired two-tailed Student’s t-test. *p < 0.05, **(D)** Confocal images show prominent SAA expression in biopsies of inflamed tissues from UC patients. Scale bar corresponds to 100µm. SAA1/2 (green), EPCAM (red) and nucleus (Draq7; blue).

**Figure S4.**
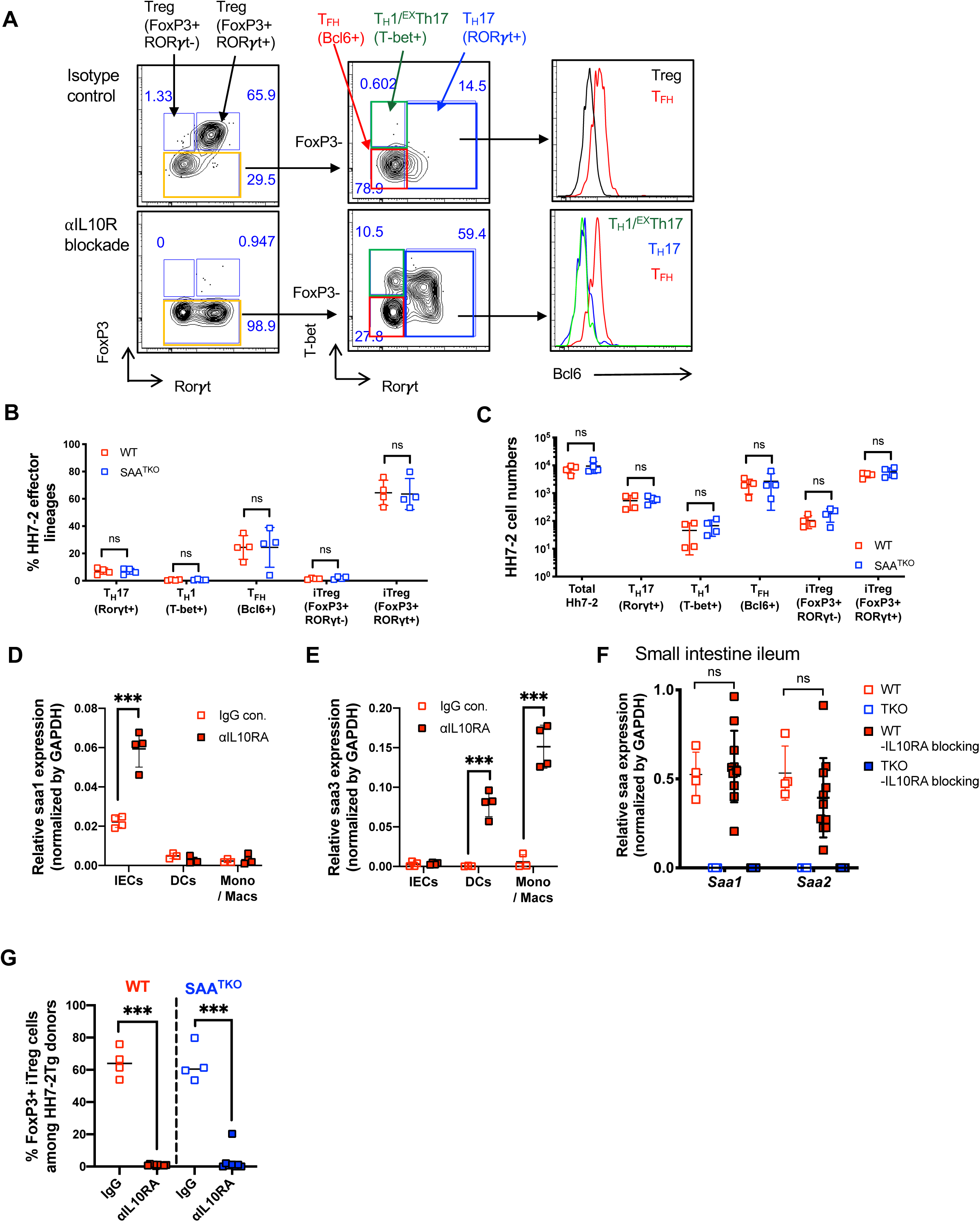
Role of SAAs in IL-10R blockade-induced colitis in response to *Helicobacter hepaticus*, Related to Figure 4. **(A)** Gating strategy to identify all T_H_ populations amongst HH7-2tg donor-derived cells in the colon lamina propria of representative homeostatic (isotype control IgG-injected) or colitis-developing (anti-IL10R-injected) recipients at 2 weeks post transfer. **(B and C)** Characterization of HH7-2tg donor cells in colon lamina propria of recipient mice injected with isotype control Ab. SAA^TKO^ (blue boxes, n = 4) and WT (red boxes, n = 4) littermates. Frequencies **(B)** and numbers **(C)** of indicated T_H_ cells based on transcription factor staining. **(D and E)** Normalized expression of *Saa1* **(D)** and *Saa3* **(E)** in the indicated sort-purified populations from colons of mice treated as in Figure 4A. Tissues were harvested 3-5 days after transfer of HH7-2tg cells. **(F)** Normalized expression of *Saa1* and *Saa2* in ileum of recipient mice. **(G)** Percentage of Foxp3^+^ iTreg cells in isotype-treated recipients (open boxes) versus αIL10RA-treated recipients (solid boxes). **(A-G)** Experiments were conducted based on the same scheme as in Figure 4A. Statistics were calculated using the two-stage step-up method of Benjamini, Krieger and Yekutieliun. Error bars denote the mean ± s.d. ns = not significant. *p<0.05, ***p < 0.001.

**Figure S5.**
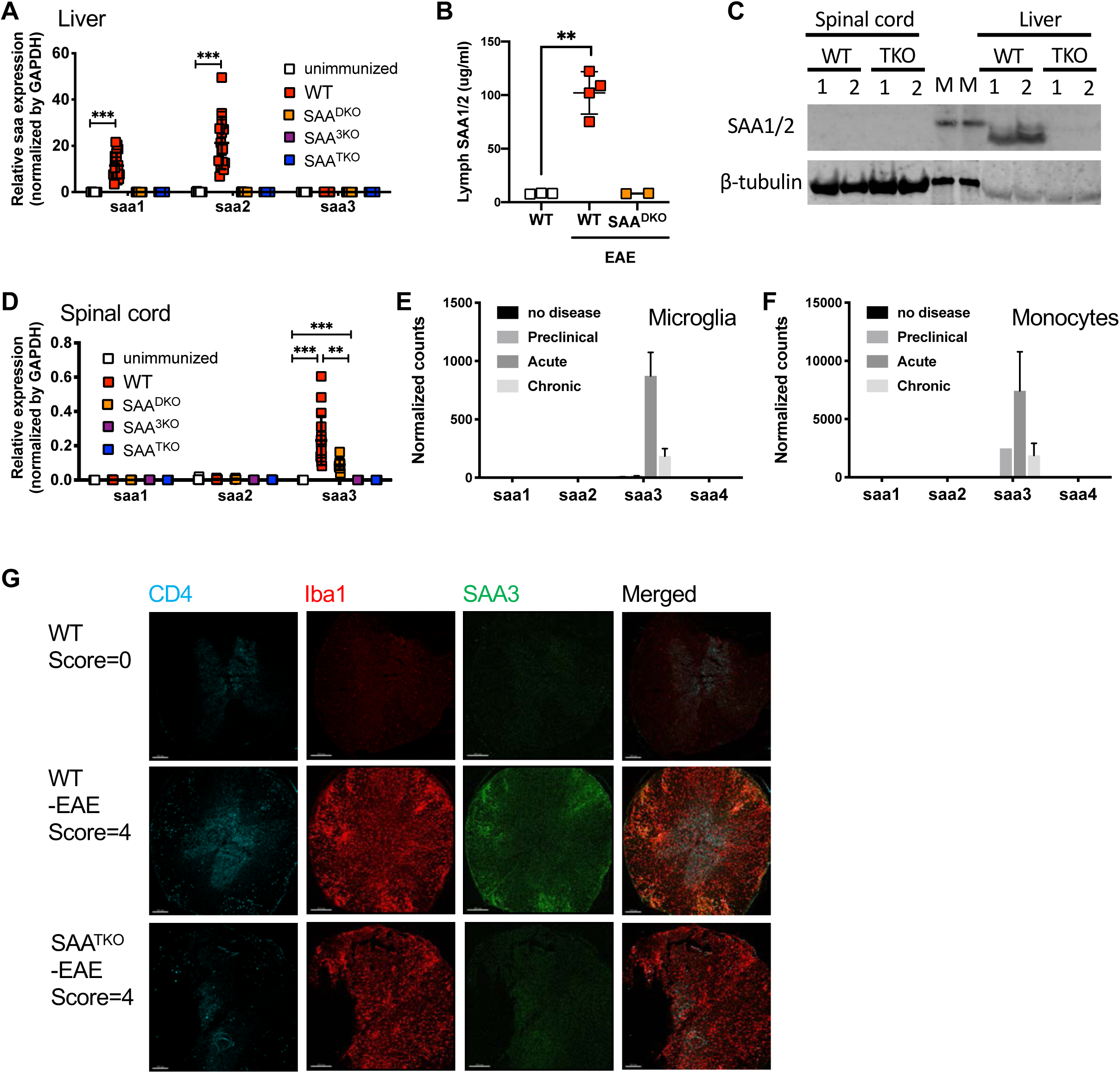
Distinct sources of SAA production in EAE, Related to Figure 5. **(A)** Normalized expression of *Saa* isotypes in the liver at day 15 post MOG-immunization. **(B)** SAA1/2 concentrations in lymph fluid collected from thoracic duct, at day 10 post immunization (pre-clinical stage) of EAE. **(C)** Expression of SAA1/2 in the spinal cord and liver of WT (n=2) and SAA^TKO^ (n=2) at day 15 of EAE. β-tubulin is shown as a loading control. Center lanes marked (M) are molecular weight markers, 14kD for the top and 64kD for the bottom. **(D)** Normalized expression of *Saa* isotypes in the spinal cord at day 15 post MOG-immunization. **(E and F)** Expression of SAAs in myeloid cells of the CNS. In order to distinguish microglia (DsRed^+^) from infiltrating monocyte-derived macrophages, *Cx3cr1^creER^:R26-DsRed* mice were injected with tamoxifen at day 28 and day 30, then rested for 30 days before MOG-immunization. RNAseq was performed on sort-purified microglia and infiltrating monocytes. Normalized RNAseq counts of *Saa* isotypes in microglia (**E**) and monocyte (**F**) isolated from CNS of WT mice at the indicated stages of EAE. Data for each condition are the mean of 2 biological replicates. **(G)** Representative confocal image of spinal cord cross sections illustrating the specificity of the α-SAA3 Ab used. IBA1 (red), SAA3 (green), and CD4 (aqua). **(A, B, and D)** Statistics were calculated using the unpaired two-sided Welch’s t-test. Error bars denote the mean ± s.d. **p < 0.01, and ***p < 0.001.

**Figure S6.**
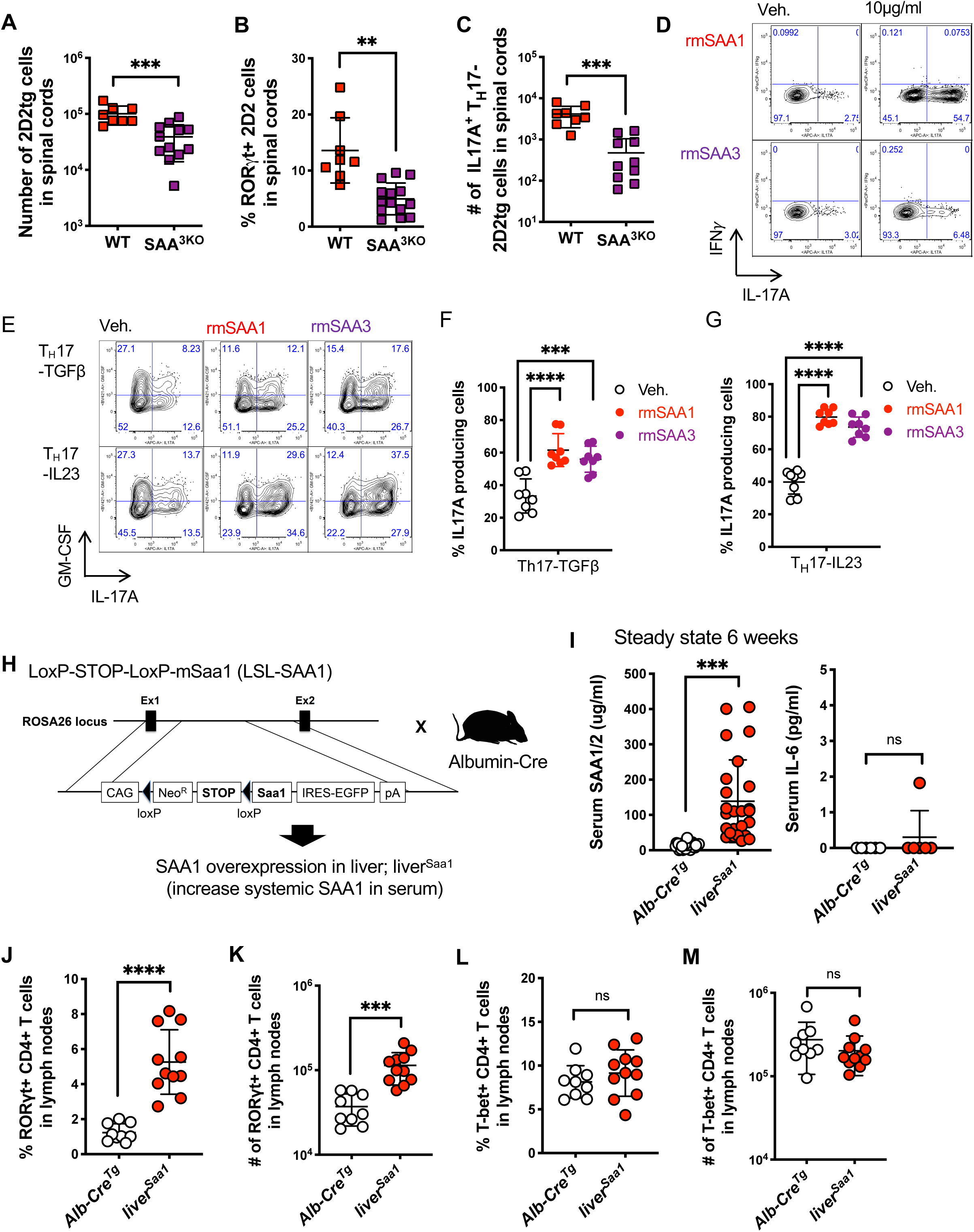
Role of SAAs in the T_H_17-dependent 2D2 transfer EAE model. **(A-C)** Total number **(A)**, percentage that are RORγt^+^ T_H_17 **(B)** and quantification of IL-17A^+^ **(C)** transferred 2D2tg donor cells in the CNS of WT (red boxes, n = 8) or SAA^3KO^ (purple boxes, n = 9) recipients on day 17 post-adoptive transfer. Summary of 3 experiments. **(D)** In vitro Th17 cell differentiation in response to SAA1 versus SAA3. The experiment was conducted based on the scheme in Figure 1A. Expression of IL-17A versus IFN-γ in *in vitro* polarized CD4^+^ T cell with IL-6 and TGFβ neutralizing antibodies ± rmSAA1 (10μg/ml) or ± rmSAA3 (10μg/ml) following restimulation. **(E-G)** Comparison of cytokine production following different Th17 cell differentiation conditions. Experiments were conducted based on the scheme in Figure 1A. **(E)** Expression of IL-17A versus GM-CSF in *in vitro* polarized CD4^+^ T cell with T_H_17-TGFβ (Top: IL6 + TGFβ) or T_H_17-IL23 (Bottom: IL-6 + IL-1β + IL-23) conditions ± rmSAA1 (10μg/ml) or ± rmSAA3 (10μg/ml) following restimulation. **(F and G)** Summary of IL-17A frequency. Summary of 4 experiments, with 8 biological replicates. **(H)** Schematic of conditional *Saa1* targeting construct and generation of liver^Saa1^mouse. **(I)** Serum concentrations of SAAs and IL-6 in liver-overexpressing mice. Left: SAA1/2 in serum of liver^Saa1^ (red circles, n = 27) or Alb-Cre^Tg^ (white circles, n = 25) and right: IL-6 liver^Saa1^ (red circles, n = 6) or Alb-Cre^Tg^ (white circles, n = 6) in 6-week-old mice. **(J and K)** Percent (**J**) and total number (**K**) of RORγt^+^ T_H_17 cells amongst Foxp3^Neg^ CD44^hi^ CD4^+^ T cells in secondary lymph nodes of 6-week-old mice with liver^Saa1^ (red circles, n = 10) or Alb-Cre^Tg^ (white circles, n = 10) summarized over two experiments. **(L and M)** As in **(J and K)**, but of Tbet^+^ T_H_1 cells. **(A-G and I-M)** Statistics were calculated using the unpaired two-sided Welch’s t-test. Error bars denote the mean ± s.d. ns = not significant, *p < 0.05, **p < 0.01, ***p < 0.001, ****p < 0.0001.

